# Breadth of SARS-CoV-2 Neutralization and Protection Induced by a Nanoparticle Vaccine

**DOI:** 10.1101/2022.01.26.477915

**Authors:** Dapeng Li, David R. Martinez, Alexandra Schäfer, Haiyan Chen, Maggie Barr, Laura L. Sutherland, Esther Lee, Robert Parks, Dieter Mielke, Whitney Edwards, Amanda Newman, Kevin W. Bock, Mahnaz Minai, Bianca M. Nagata, Matthew Gagne, Daniel C. Douek, C. Todd DeMarco, Thomas N. Denny, Thomas H. Oguin, Alecia Brown, Wes Rountree, Yunfei Wang, Katayoun Mansouri, Robert J. Edwards, Guido Ferrari, Gregory D. Sempowski, Amanda Eaton, Juanjie Tang, Derek W. Cain, Sampa Santra, Norbert Pardi, Drew Weissman, Mark A. Tomai, Christopher B. Fox, Ian N. Moore, Hanne Andersen, Mark G. Lewis, Hana Golding, Robert Seder, Surender Khurana, Ralph S. Baric, David C. Montefiori, Kevin O. Saunders, Barton F. Haynes

## Abstract

Coronavirus vaccines that are highly effective against SARS-CoV-2 variants are needed to control the current pandemic. We previously reported a receptor-binding domain (RBD) sortase A-conjugated ferritin nanoparticle (RBD-scNP) vaccine that induced neutralizing antibodies against SARS-CoV-2 and pre-emergent sarbecoviruses and protected monkeys from SARS-CoV-2 WA-1 infection. Here, we demonstrate SARS-CoV-2 RBD-scNP immunization induces potent neutralizing antibodies in non-human primates (NHPs) against all eight SARS-CoV-2 variants tested including the Beta, Delta, and Omicron variants. The Omicron variant was neutralized by RBD-scNP-induced serum antibodies with a mean of 10.6-fold reduction of ID50 titers compared to SARS-CoV-2 D614G. Immunization with RBD-scNPs protected NHPs from SARS-CoV-2 WA-1, Beta, and Delta variant challenge, and protected mice from challenges of SARS-CoV-2 Beta variant and two other heterologous sarbecoviruses. These results demonstrate the ability of RBD-scNPs to induce broad neutralization of SARS-CoV-2 variants and to protect NHPs and mice from multiple different SARS-related viruses. Such a vaccine could provide the needed immunity to slow the spread of and reduce disease caused by SARS-CoV-2 variants such as Delta and Omicron.

## INTRODUCTION

Despite the remarkable success of approved COVID-19 vaccines, additional broadly protective vaccines may be needed to combat breakthrough infections caused by emerging SARS-CoV-2 variants and waning immunity. Moreover, pan-Sarbecovirus vaccines are needed for the prevention of new animal SARS-like viruses that may jump to humans in the future (Levin et al., 2021). Modified mRNA vaccines encapsulated in lipid nanoparticles (LNPs) have proved transformative for COVID-19 vaccine development and for vaccine development in general (Chaudhary et al., 2021; Pardi et al., 2018; Pardi et al., 2020). Developed in 11 months and providing >90% efficacy from transmission, the mRNA-1273 and the BNT162b2 COVID-1 vaccines, while showing the most reduction in efficacy from SARS-CoV-2 Beta and Omicron variants, continue to provide significant protection from serious COVID-19 disease, hospitalization, and death (Baden et al., 2021; Polack et al., 2020). The Omicron variant, however, has proved to be more transmissible than previous variants, now accounting for the majority of global isolates (http://www.gisaid.org/hcov19-variants). Likely arising from immunocompromised individuals in South Africa, the Omicron variant spike protein contains 30 mutations compared to the WA-1 strain, and continues to evolve (Wang and Cheng, 2021). While likely less pathogenic than Delta and other SARS-CoV-2 variants, the enhanced transmissibility of Omicron, coupled with the sheer number of resulting cases, has resulted in a higher absolute number of COVID-19 patients compared to previous variant infections, thus providing a continued burden on global health care systems.

We previously reported an RBD-based, sortase A-conjugated nanoparticle (RBD-scNP) vaccine formulated with the TLR7/8 agonist 3M-052-aqueous formulation (AF) (hereafter 3M-052-AF) plus Alum, that elicited cross-neutralizing antibody responses against SARS-CoV-2 and other sarbecoviruses, and protected against the WA-1 SARS-CoV-2 strain in non-human primates (NHPs) (Saunders et al., 2021). Here, we found RBD-scNPs induced antibodies that neutralized all variants tested including Beta and Omicron, and protected against Beta and Delta variant challenges in macaques. Moreover, RBD-scNP immunization protected in highly susceptible aged mouse models against challenges of SARS-CoV- 2 Beta variant and other sarbecoviruses. In addition, while formulating RBD-scNP with Alum, 3M-052-AF, or 3M-052-AF + Alum each protected animals from WA-1 challenge, the 3M-052-AF/ RBD-scNP formulation was optimal for induction of neutralization titers to variants and protection from lung inflammation. Finally, we found that RBD-, N-terminal domain (NTD)- and spike-2P (S2P)-scNPs each protected comparably in the upper and lower airways from WA-1, but boosting with the NTD-scNP protected less well than RBD-scNP or S2P-scNP.

## RESULTS

### RBD-scNPs induce neutralizing antibodies against SARS-CoV-2 B.1.1.529 (Omicron) and other variants

RBD-scNPs were used to immunize macaques 3 times four weeks apart (***Figure 1A***). To test whether RBD-scNP-induced antibodies can neutralize SARS-CoV-2 variants, we collected macaque plasma samples two weeks after the 3^rd^ RBD-scNP immunization (Saunders *et al*., 2021) and assessed their ability to neutralize pseudovirus infection of 293T-ACE2-TMPRSS2 cells by SARS-CoV-2 WA-1 and 8 variants (***Figure 1A***). RBD-scNP induced potent plasma neutralizing antibodies against the WA-1 strain with geometric mean titer (GMT) ID50 of 12,266.7, while reduced ID50 titers were observed to different extents for the variants (***Figure 1B-C***). In particular, the highest reduction of neutralizing activity was observed for the B.1.351 (Beta) variant, which ranged from 4.1- to 10.2- fold (***Figure 1C***). To determine if the B.1.1.529 (Omicron) variant could escape RBD-scNP-induced neutralizing antibodies, we compared immune sera for capacity to neutralize the D614G and Omicron variants in a 293T-ACE2 pseudovirus assay. Serum antibodies induced by three doses of RBD-scNP immunization (Saunders *et al*., 2021) neutralized both the D614G (GMT ID50=40,249, GMT ID80=12,053) and Omicron (GMT ID50=3,794, GMT ID80=864) pseudoviruses. A 10.6-fold drop in the GMT ID50 and 14.0-fold drop in the GMT ID80 were observed (***Figure 1D***). Thus, high titers of neutralizing antibodies against SARS-CoV-2 variants, including Omicron, were elicited by the RBD-scNP immunization in macaques.

**Figure 1.**
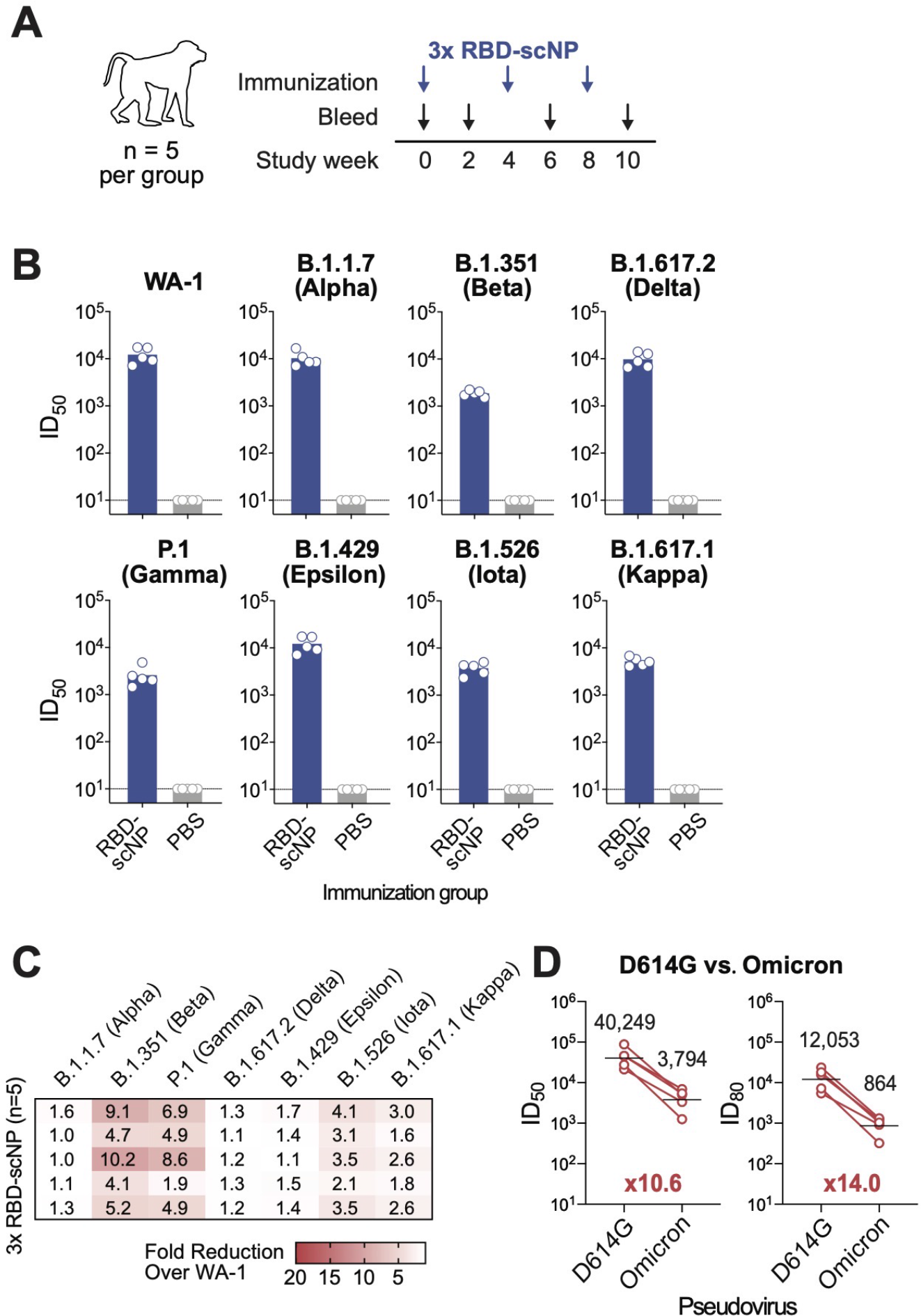
RBD-scNP vaccination elicits broad neutralizing antibodies against SARS-CoV-2 variants in macaques. **(A)** Schematic of the vaccination study. Cynomolgus macaques (*n*=5 per group) were immunized 3 times with PBS control or 100 μg RBD-scNP adjuvanted with 3M-052-AF + Alum. **(B-C)** Plasma antibody (post-3^rd^ immunization) neutralization of SARS-CoV-2 variants pseudovirus infection in 293T-ACE2-TMPRSS2 cells. (B) Neutralization 50% inhibitory dilution (ID_50_) titers. Each symbol represents an individual macaque. Bars indicate group geometric mean ID_50_. (C) Reduction of ID_50_ titers against variants were shown as fold reduction compared to the titers against WA-1. Each row shows values for an individual macaque for each virus. **(D)** Plasma antibody (post-3^rd^ immunization) neutralization titers against pseudoviruses of the SARS-CoV-2 Omicron variants in 293T-ACE2 cells. The geometric mean ID_50_ and ID_80_ titers and the fold reduction compared to D614G are shown.

### RBD-scNPs protect macaques from SARS-CoV-2 WA-1, Beta and Delta challenge

Next, we determined if two doses of RBD-scNP vaccination could protect NHPs from challenge by SARS-CoV-2 WA-1, Beta or Delta variants. We immunized cynomolgus macaques with two doses of RBD-scNP vaccine, PBS, or adjuvant alone (***Figure 2A***). We also immunized a group of macaques with soluble RBD as a comparator to RBD-scNP immunization to test the effects of multimerization. RBD-scNP and soluble RBD monomer immunization elicited similar titers of antibodies binding to SARS-CoV-2 and other CoV spike antigens (***Supplementary Figure 1A***), which also similarly blocked ACE2- binding on SARS-CoV-2 spike and bat CoV RsSHC014 spike (***Supplementary Figure 1B-C***). RBD- scNPs and soluble RBD induced similar levels of antibodies targeting the sarbecovirus cross-neutralizing DH0147-epitope (Hastie et al., 2021; Li et al., 2021a; Li et al., 2021b; Martinez et al., 2021a) on SARS-CoV-2 spike as well as on RsSHC014 spike (***Supplementary Figure 1B-C***). In pseudovirus neutralization assays, the RBD-scNP group exhibited higher titers of neutralizing antibodies than the soluble RBD group against the WA-1, Alpha, Epsilon, Iota, and Delta viruses, with comparable neutralizing titers against Beta, Gamma, and Kappa variants (***Figure 2B***). In both groups, neutralizing titers for the Beta, Gamma, and Iota variants were reduced compared to WA-1 (***Supplementary Figure 1D***). In addition, serum antibodies induced by two doses of RBD-scNP immunization exhibited modest neutralization against Omicron pseudovirus (GMT ID50=880, GMT ID80=280), with a 63-fold drop in the GMT ID50 and 37-fold drop in the GMT ID80 compared to D614G neutralization titers (***Figure 2C***). Thus, RBD-scNP induced higher neutralizing antibodies than soluble RBD monomer for 5 out of 8 SARS-CoV-2 variant pseudoviruses tested. RBD-scNP showed reduction of Omicron neutralization titers compared to the D614G variant after two immunizations, with far less fold-reduction in Omicron neutralization titers after three immunizations (***Figure 1D***).

**Figure 2.**
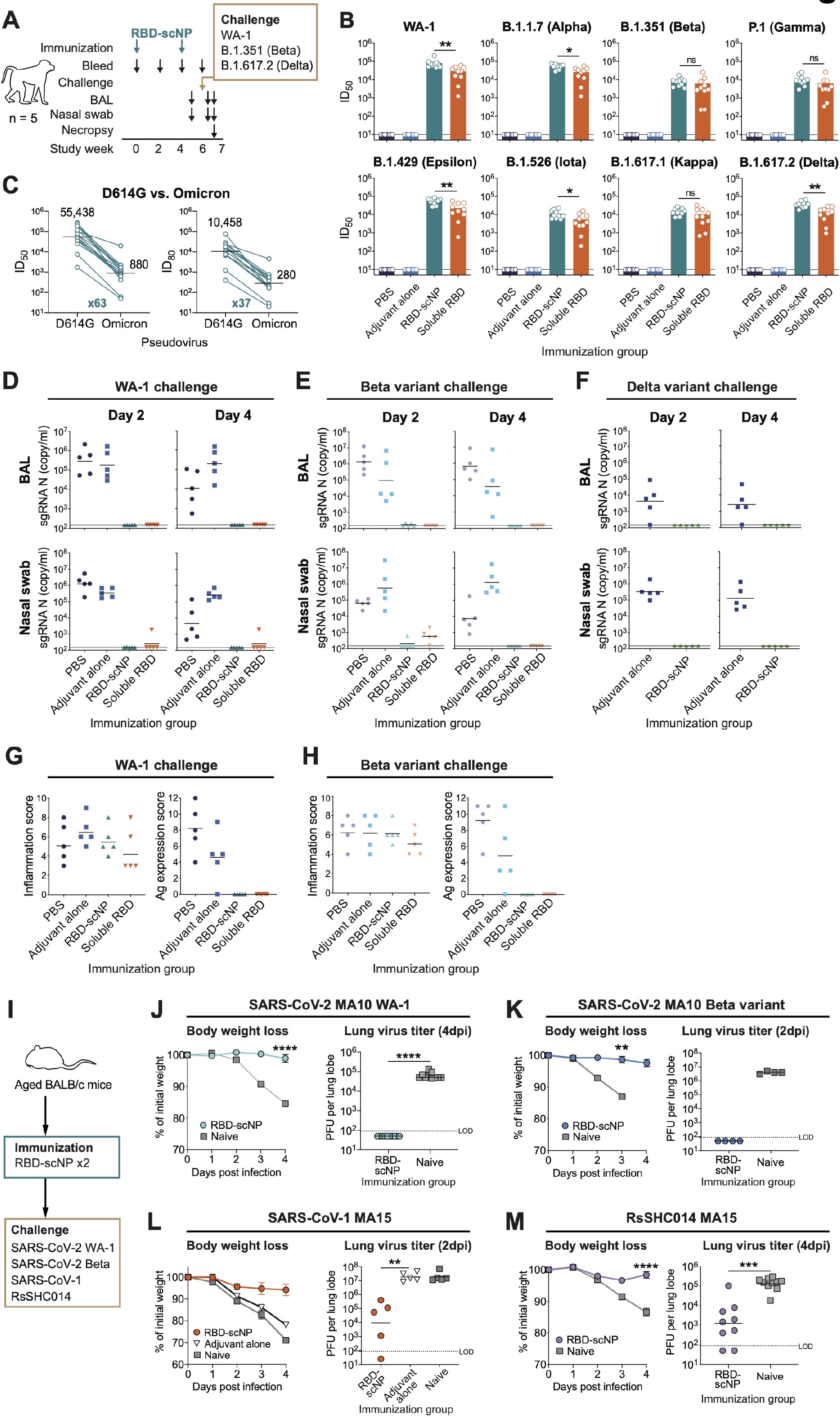
Two doses of RBD-scNP vaccination protected non-human primates and mice from challenges of SARS-CoV-2 variants and other betacoronaviruses. **(A)** Schematic of the vaccination and challenge studies. Cynomolgus macaques were immunized twice and challenged with SARS-CoV-2 WA-1 strain (*n*=5) or B.1.351 (Beta; *n*=5) or B.1.617.2 (Delta; *n*=5). Bronchoalveolar lavage (BAL) and nasal swab samples were collected for subgenomic (sgRNA) viral replication tests. Animals were necropsied on day 4 post-challenge for pathologic analysis. **(B)** Plasma antibody neutralization ID_50_ titers against pseudoviruses of SARS-CoV-2 variants in 293T-ACE2-TMPRSS2 cells. **(C)** Plasma antibody neutralization ID_50_ titers against pseudoviruses of SARS-CoV-2 Omicron variants in 293T-ACE2 cells. The geometric mean ID_50_ and ID_80_ titers and the fold reduction compared to D614G are shown. **(D-F)** SARS-CoV-2 sgRNA levels for nucleocapsid (N) gene in BAL and nasal swab samples collected on day 2 and 4 after SARS-CoV-2 WA-1 (D), Beta variant (E) or Delta variant (F) challenge. Dashed line indicates limit of the detection (LOD). **(G-H)** Histopathological analysis of the SARS-CoV-2 WA-1 (G) and Beta variant (H) challenged monkeys. Scores of lung inflammation determined by haematoxylin and eosin (H&E) staining and SARS-CoV-2 nucleocapsid antigen (Ag) expression determined by immunohistochemistry (IHC) staining. **(I)** Schematic of the mouse challenge studies. Aged female BALB/c mice (n=10 per group) were immunized intramuscularly twice and challenged with SARS-CoV-2 mouse-adapted 10 (MA10) WA-1, SARS-CoV-2 MA10 Beta variant, SARS-CoV-1 mouse-adapted 15 (MA15), or Bat coronavirus (CoV) RsSHC014 MA15. **(J)** Weight loss and lung virus titers at 4 days post-infection (dpi) of the SARS-CoV-2 MA10 WA-1 challenged mice. **(K)** Weight loss and lung virus titers at 2 dpi of the SARS-CoV-2 MA10 Beta variant challenged mice. **(L)** Weight loss and lung virus titers at 2 dpi of the SARS-CoV-1 MA15 challenged mice. **(M)** Weight loss and lung virus titers at 4 dpi of the Bat CoV RsSHC014 MA15 challenged mice. ns, not significant, *P<0.05, **P<0.01, ***P<0.001, ****P<0.0001, Wilcoxon rank sum exact test.

Two weeks after the second vaccination, macaques (n=5 per group) were challenged with SARS-CoV-2 WA-1, SARS-CoV-2 B.1.351 (Beta) variant, or SARS-CoV-2 B.1.617.2 (Delta) variant (***Figure 2A***). In the PBS or adjuvant alone group, high copies of envelope (E) and nucleocapsid (N) gene subgenomic RNA (sgRNA) were detected in both Bronchoalveolar Lavage (BAL) and nasal swab samples collected on day 2 and 4 post-challenge (***Figure 2D-F and Supplementary Figure 1E-G***). In contrast, 5 of 5 animals in the RBD-scNP group, and 4 of 5 animals in the soluble RBD group, were completely protected from WA-1 infection, as indicated by no detectable sgRNA in either BAL or nasal swab (***Figure 2D***). After the SARS-CoV-2 Beta variant challenge, nasal N gene sgRNA was detected in only 1 of 5 of the RBD-scNP immunized monkeys but in 4 of 5 of the soluble RBD immunized monkeys (***Figure 2E***). In addition, after the SARS-CoV-2 Delta variant challenge, all animals that received two doses of RBD-scNP immunization showed no detectable sgRNA in BAL or nasal swab samples (***Figure 2F***).

Animals were necropsied 4 days after challenge for histopathologic analysis to determine SARS-CoV-2-associated lung inflammation. After WA-1 and Beta variant challenge, lung tissue haematoxylin and eosin (H&E) staining revealed no difference between groups (***Figure 2G-H***). However, immunohistochemistry (IHC) staining showed the presence of SARS-CoV-2 nucleocapsid antigen in the lungs of macaques administered PBS or adjuvant alone, but not in the lungs of RBD-scNP or soluble RBD immunized monkeys (***Figure 2G-H***). Thus, while lung inflammation was observed in immunized macaques, two doses of RBD-scNP immunization protected against viral replication of WA-1, the Beta variant, or Delta variant in both lower and upper airways. In addition, RBD-scNP was superior to soluble RBD in terms of protecting from the hard-to-neutralize Beta variant infection in the upper respiratory tract.

### RBD-scNPs induce protective responses in mice against SARS-CoV-2 Beta variant and other sarbecoviruses

To define the protective efficacy of the RBD-scNP vaccination against different sarbecoviruses, we immunized aged mice with two doses of RBD-scNPs, challenged the mice with mouse-adapted SARS-CoV-2 WA-1, SARS-CoV-2 Beta variant, SARS-CoV-1, or bat CoV RsSHC014 (***Figure 2I***). In the SARS-CoV-2 WA-1 challenge study, RBD-scNP protected mice from weight loss through 4 days post infection (dpi) and protected from viral replication in the lungs (***Figure 2J***). Similar protection from weight loss and lung viral replication were observed in the SARS-CoV-2 Beta variant challenged mice (***Figure 2K***); moreover, by 4 dpi mortality was observed in the unimmunized mice group but no RBD-scNP-immunized mice died. Mice immunized with RBD-scNP were also protected against weight loss induced by SARS-CoV-1 and showed ∼3-log lower average PFU titer in lungs compared to adjuvant alone and unimmunized groups (***Figure 2L***). Lastly, RBD-scNP immunization conferred protection against RsSHC014 challenge-induced weight loss and resulted in ∼2-log lower average PFU titer than naïve mice (***Figure 2M***). Thus, two doses of RBD-scNP immunization elicited protective immune responses against SARS-CoV-2 Beta variant and multiple other sarbecoviruses in aged mouse models.

### Adjuvant is required for RBD-scNP induction of potent plasma and mucosal antibody responses

To optimize adjuvant formulations for the RBD-scNP vaccine, we next formulated the RBD-scNP immunogen with the TLR7/8 agonist 3M-052-AF alone, with aluminum hydroxide (Alum) alone, or with 3M-052-AF adsorbed to Alum (3M-052-AF + Alum). Control groups included NHPs immunized with immunogen alone (RBD-scNP without adjuvant), adjuvant alone (3M-052-AF, Alum, or 3M-052-AF + Alum without immunogen), or PBS alone (***Figure 3A***). After three immunizations, RBD-scNP alone without adjuvant induced minimal binding antibodies to SARS-CoV-2 and other CoV spike antigens, whereas higher titers of binding antibodies were induced by RBD-scNP formulated with each adjuvant (***Supplementary Figure 2A***). While all three adjuvant formulations were highly immunogenic, RBD-scNP adjuvanted with 3M-052-AF induced the highest DH1047-blocking plasma antibodies (*p<*0.05; Wilcoxon rank sum exact test; ***Supplementary Figure 2B-C***). Mucosal antibody levels tended to be comparable for macaques who received RBD-scNP formulated with 3M-052-AF or 3M-052+Alum, with only low titers being seen when Alum was used to adjuvant the RBD-scNP (***Supplementary Figure 2D- E***).

**Figure 3.**
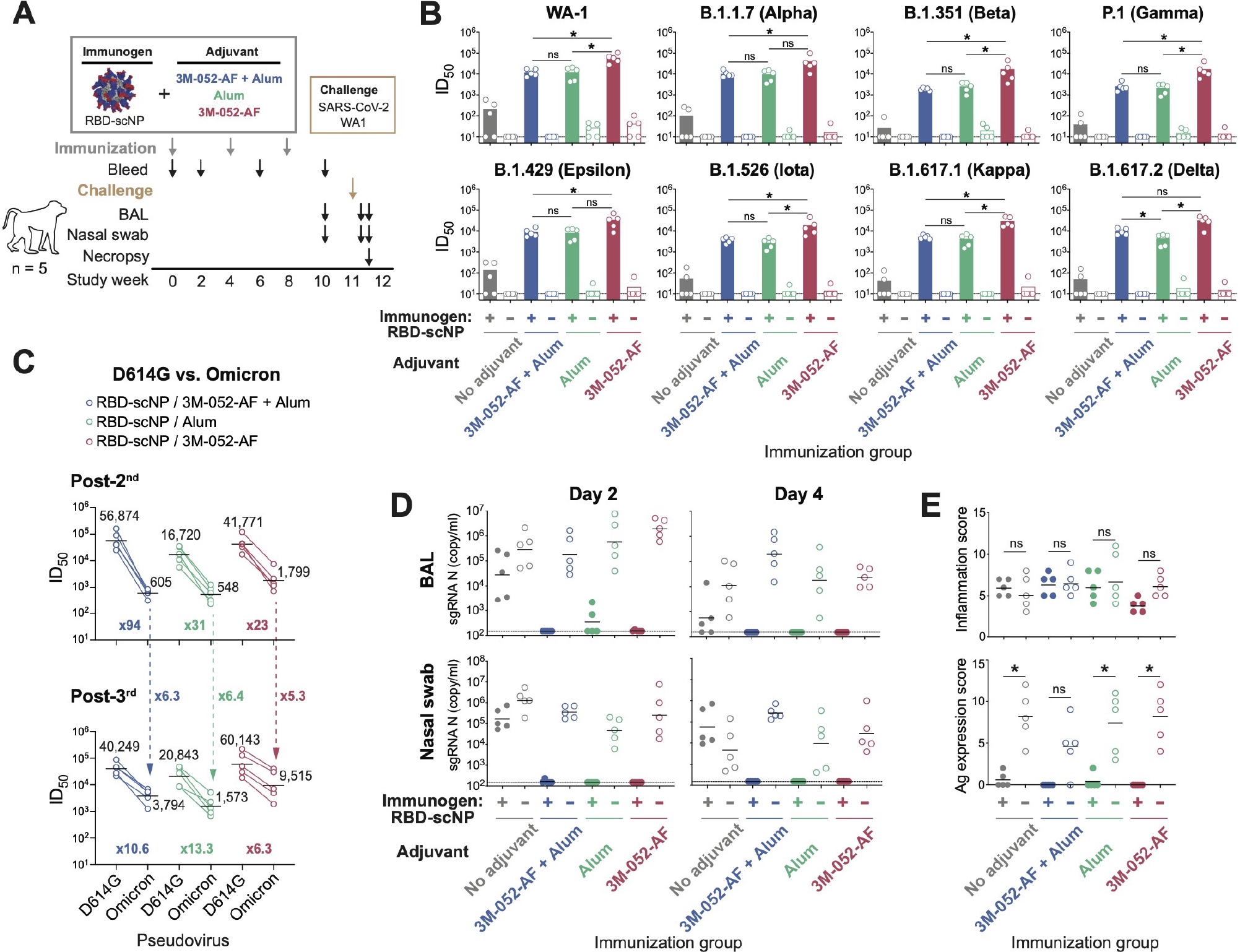
Neutralizing antibodies and *in vivo* protection elicited by RBD-scNP vaccine formulated with three different adjuvants. **(A)** Schematic of the vaccination and challenge study. Cynomolgus macaques (*n*=5 per group) were immunized intramuscularly 3 times with 100 μg of RBD-scNP adjuvanted with 3M052-AF + Alum, Alum, 3M052-AF, or PBS control. Animals injected with adjuvant alone or PBS were set as control groups. Monkeys were then challenged with SARS-CoV-2 WA-1, collected for blood, BAL and nasal swab samples, and necropsied for pathologic analysis. **(B)** Neutralization ID_50_ titers of plasma antibodies (post-3^rd^ immunization) against pseudovirus of SARS-CoV-2 variants in 293T-ACE2-TMPRSS2. **(C)** Plasma antibody (post-2^nd^ and post-3^rd^ immunization) neutralization titers against pseudoviruses of the SARS-CoV-2 Omicron variants in 293T-ACE2 cells. The geometric mean ID50 titers and the fold reduction compared to D614G are shown. The dashed arrows indicate fold increase of ID_50_ titer induced by the 3^rd^ boost. **(D)** SARS-CoV-2 N gene sgRNA in BAL and nasal swab samples collected on day 2 and 4 post-challenge. Dashed line indicates limit of the detection. **(E)** Histopathological analysis. Lung sections from each animal were scored for lung inflammation by H&E staining, and for SARS-CoV-2 nucleocapsid Ag expression by IHC staining. ns, not significant, *P<0.05, Wilcoxon rank sum exact test.

### Robust neutralizing antibodies and *in vivo* protection induced by adjuvanted RBD-scNP

While the RBD-scNP alone group showed minimal neutralizing antibody titers, the RBD-scNP + 3M-052-AF group had remarkably high pseudovirus neutralizing antibody titers against SARS-CoV-2 WA-1 strain (GMT ID50 = 59,497). The GMT ID50 of RBD-scNP + 3M-052-AF + Alum and RBD-scNP + Alum groups against WA-1 were 12,267 and 12,610, respectively (***Figure 3B***). Moreover, RBD-scNPs + 3M-052-AF immunized animals exhibited the highest magnitudes of neutralizing antibodies against each variant we tested (***Figure 3B***). Serum antibodies induced by two immunizations of RBD-scNP showed 94-, 31- and 23-fold drop in the Omicron ID50 GMT compared to D614G neutralization titers for 3M-052-AF + Alum, Alum, and 3M-052-AF adjuvant groups respectively (***Figure 3C***, upper panel), whereas after three immunizations the reduction-fold of Omicron ID50 GMT were 10.6-, 13.3-, and 6.3- fold in the 3M-052-AF + Alum, Alum, and 3M-052-AF adjuvated groups, respectively (***Figure 3C,*** lower panel). The most potent neutralizing antibodies against Omicron were observed in the 3M-052-AF-adjuvanted group, showing ID50 GMTs of 1,799 after the 2^nd^ dose and 9,515 after the 3^rd^ dose (***Figure 3C***). Importantly, although the 3^rd^ boost occurred only 1 month after the 2^nd^ boost, it resulted in a 5.3-fold increase in Omicron titer with 3M-052-AF adjuvant, 6.4-fold increase with Alum, and 6.3-fold increase with 3M-052-AF + Alum (***Figure 3C***). Thus, a third boost after only 1 month was effective in enhancing neutralization breadth to the Omicron variant. Moreover, the 3M-052-AF adjuvant induced higher Omicron titers than 3M-052-AF + Alum or Alum alone.

To compare *in vivo* protection of RBD-scNP with different adjuvant formulations, cynomolgus macaques were challenged with the WA-1 strain of SARS-CoV-2 three weeks after the third immunization (***Figure 3A***). Compared to unimmunized monkeys, the adjuvant alone groups exhibited similar or higher levels of E and N sgRNA, and the RBD-scNP immunogen alone reduced sgRNA copies by only ∼1-2 logs (***Figure 3D and Supplementary Figure 2F***). Immunization with either RBD-scNP adjuvanted with 3M-052-AF + Alum or 3M-052-AF conferred robust protection against SARS-CoV-2 infection. Macaques in these groups had under-detection-limit or near-baseline levels of sgRNA N and E in both lower and upper respiratory tracts, demonstrating that adjuvant was required for eliciting potent protection from SARS-CoV-2 challenge. RBD-scNP + Alum immunized macaques showed positive E or N gene sgRNA in 1 of 5 and 2 of 5 macaques in BAL samples collected on day 2 post-challenge respectively. By day 4 post-challenge, all RBD-scNP adjuvanted groups showed no detectable sgRNA (***Figure 3D and Supplementary Figure 2F***).

Histologic analysis of lung tissue showed that RBD-scNP + adjuvant and adjuvant only groups had similar inflammation scores (***Figure 3E***). IHC staining of the lung tissues exhibited high SARS-CoV-2 nucleocapsid antigen expression in the unimmunized and adjuvant alone groups. In contrast, 1 of 5 of the RBD-scNP + Alum immunized animals and 2 of 5 of the immunogen alone immunized animals had low level nucleocapsid antigen expression, and no viral antigen was detected in the RBD-scNP plus 3M-052- AF + Alum or RBD-scNP plus 3M-052-AF immunized animals (***Figure 3E***). Therefore, the three adjuvants conferred comparable protection against viral replication by day 4 post-challenge, with 3M- 052-AF and 3M-052-AF + Alum providing the best reduction in virus replication when adjuvanting RBD-scNP.

### RBD-scNP, NTD-scNP, and S2P-scNP vaccines induce both neutralizing and ADCC-mediating antibodies

While 90% of neutralizing antibodies target the RBD, neutralizing antibodies can target other sites on spike. Thus, we generated scNPs with NTD and S-2P and compared the antibody response elicited by these ferritin nanoparticles to RBD-scNPs (***Figure 4A-B and Supplementary Figure 3A-C***). Cynomolgus macaques were immunized three times with one of the scNPs formulated with 3M-052-AF + Alum. After three immunizations, Spike binding, ACE2-blocking, and neutralizing antibody-blocking antibodies were observed in all three groups (***Supplementary Figure 3D-F***). In the RBD-scNP and S2P-scNP immunized animals, neutralizing antibodies against SARS-CoV-2 D614G pseudovirus were detected after the first dose and were boosted after the second and third dose at week 6 and 10 (***Figure 4C***). The NTD-scNP induced sera contained IgGs that blocked NTD neutralizing antibody DH1050.1 binding (***Supplementary Figure 3E***). However, NTD-scNP-sera post 2^nd^ and 3^rd^ immunization failed to neutralize the SARS-CoV- 2 pseudovirus (***Figure 4C***) but neutralized live SARS-CoV-2 WA-1 virus in a microneutralization (MN) assay (GMT ID50=2,189) (***Figure 4D***). Importantly, S2P-scNP induced comparable plasma neutralizing antibody titers compared to RBD-scNP against SARS-CoV-2 WA-1 strain and all eight variants tested (*p*>0.05; Wilcoxon rank sum exact test; ***Figure 4E***). Among the different variants, the Beta variant showed the largest reduction in neutralization ID50 titer (5.0- to 10.9-fold) (***Figure 4F***). S2P-scNP induced neutralizing antibodies against Omicron pseudovirus, with GMT ID50 of 436 post-2^nd^ immunization (38-fold reduction) and GMT ID50 of 2,754 post-3^rd^ immunization (7.2-fold reduction) (***Figure 4G***).

**Figure 4.**
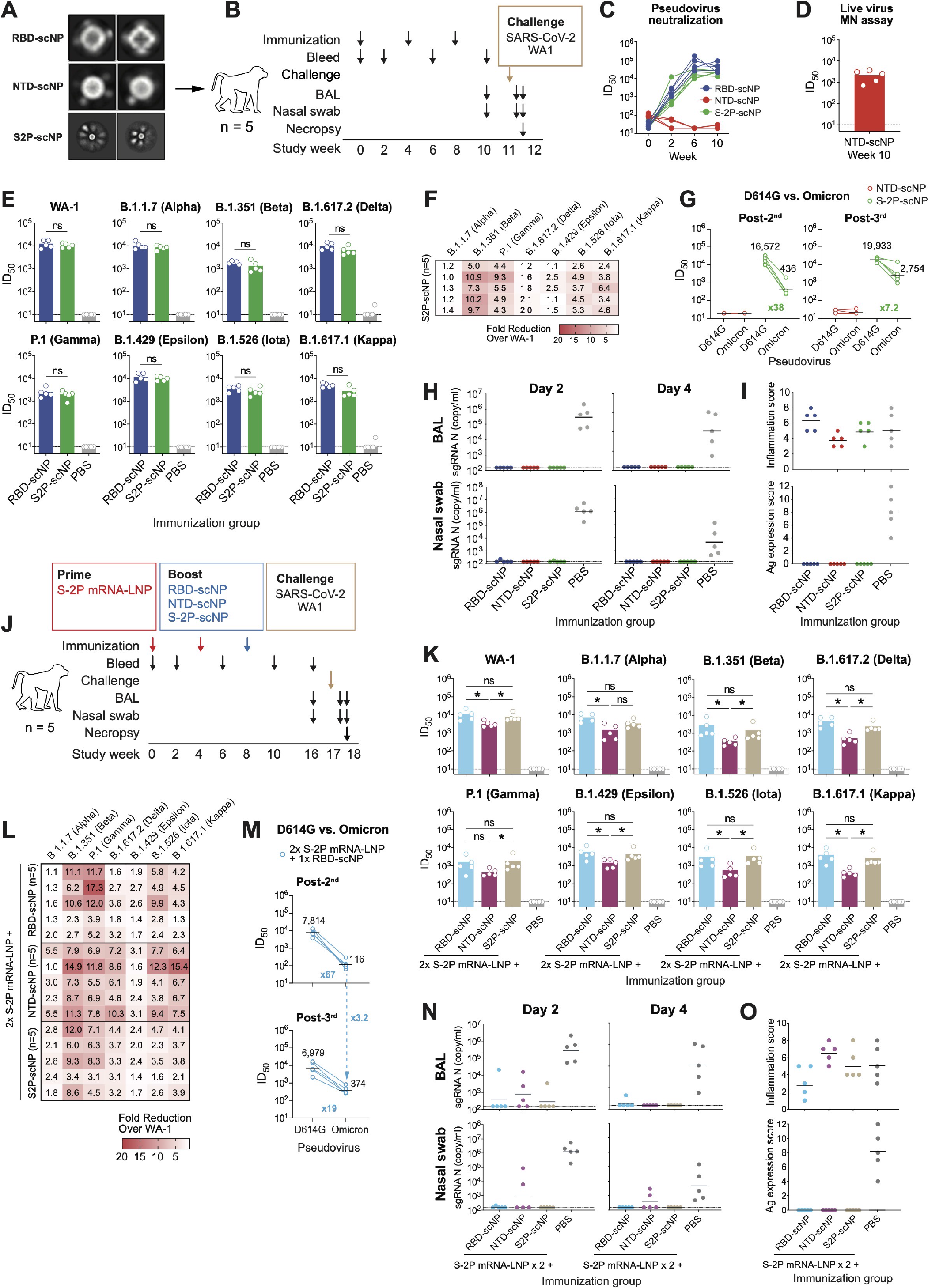
Neutralizing antibodies and *in vivo* protection induced by RBD-scNP, NTD- scNP and S2P-scNP vaccines as a three-dose regimen or as a heterologous boost for S2P mRNA-LNP vaccine. **(A)** Negative-stain electron microscopy 2D class averaging of RBD-scNP, NTD-scNP, and S2P-scNP. The size of each box: RBD-scNP and NTD-scNP, 257 Å; S2P-scNP, 1,029 Å. **(B)** Schematic of the three-dose regimen. Cynomolgus macaques (*n*=5 per group) were immunized 3 times with RBD-scNP, NTD-scNP, or S2P-scNP adjuvanted with 3M-052-AF + Alum. Monkeys were then challenged with SARS-CoV-2 WA-1, collected for blood, BAL and nasal swab samples, and necropsied for pathologic analysis. **(C)** Neutralization ID_50_ of plasma antibodies (week 0, 2, 6 and 10) against pseudotyped SARS-CoV-2 D614G strain in 293T/ACE2.MF cells. **(D)** Neutralization ID_50_ of the NTD-scNP-induced antibodies (week 10) against live SARS-CoV- 2 WA-1 virus in Vero-E6 cells in microneutralization (MN) assay. **(E-F)** Plasma antibody neutralization against pseudoviruses of the SARS-CoV-2 variants in 293T-ACE2-TMPRSS2 cells. (E) ID_50_ titers and (F) reduction of ID_50_ titers against variants were shown as fold reduction compared to the titers against WA-1. **(G)** NTD-scNP- or S2P-scNP-induced plasma antibody (post-2^nd^ and post-3^rd^ immunization) neutralization titers against pseudoviruses of the SARS-CoV-2 Omicron variants in 293T-ACE2 cells. The geometric mean ID_50_ titers and the fold reduction compared to D614G are shown. **(H)** SARS-CoV-2 N gene sgRNA in BAL and nasal swab samples collected on day 2 and 4 post-challenge. **(I)** Histopathological analysis. Scores of lung inflammation determined by H&E staining and SARS-CoV-2 nucleocapsid antigen expression determined by IHC staining. **(J)** Schematic of the heterologous prime-boost regimen. Cynomolgus macaques (*n*=5 per group) were immunized 2 times with S2P mRNA-LNP, and boosted with adjuvanted RBD-scNP, NTD-scNP, or S2P-scNP vaccine. Monkeys were then challenged with SARS-CoV-2 WA-1, collected for blood, BAL and nasal swab samples, and necropsied for pathologic analysis. **(K-L)** Plasma antibody neutralization against pseudoviruses of SARS-CoV-2 variants in 293T- ACE2-TMPRSS2 cells. (K) ID_50_ titers. (L) Reduction of ID_50_ titers against variants were shown as fold reduction compared to the titers against WA-1. **(M)** Plasma antibody (post-2^nd^ and post-3^rd^ immunization) neutralization titers against pseudoviruses of the SARS-CoV-2 Omicron variants in 293T-ACE2 cells. The geometric mean ID_50_ titers and the fold reduction compared to D614G are shown. The dashed arrow indicates fold increase of ID_50_ titer induced by the 3^rd^ boost. **(N)** SARS-CoV-2 N gene sgRNA in BAL and nasal swab samples collected on day 2 and 4 post-challenge. **(O)** Histopathological analysis. Scores of lung inflammation determined by H&E staining and SARS-CoV-2 nucleocapsid antigen expression determined by IHC staining. ns, not significant, *P<0.05, Wilcoxon rank sum exact test.

To examine other antibody functions, we examined plasma antibody binding to cell surface-expressed SARS-CoV-2 spike and antibody-dependent cellular cytotoxicity (ADCC). Plasma antibodies induced by three doses of RBD-scNP, NTD-scNP and S2P-scNP vaccination bound to SARS-CoV-2 spike on the surface of transfected cells (***Supplementary Figure 3G*)**. In a CD107a degranulation-based ADCC assay (***Supplementary Figure 3H***), plasma antibodies from all three scNP groups mediated CD107a degranulation of human NK cells in the presence of both SARS-CoV-2 spike-transfected cells and SARS-CoV-2-infected cells (***Supplementary Figure 3I***). Thus, all three scNP vaccines induced antibodies that neutralize SARS-CoV-2 and mediated ADCC.

### RBD-scNP, NTD-scNP, and S2P-scNP vaccines protect macaques against SARS-CoV-2 WA-1 challenge

To determine NTD-scNP and S2P-scNP immunization conferred protection against SARS-CoV-2, we challenged the immunized macaques with SARS-CoV-2 WA-1 strain via the intratracheal and intranasal routes after the 3^rd^ vaccination. Remarkably, all macaques that received RBD-scNP, NTD-scNP or S2P-scNP were fully protected, showing undetectable or near-detection-limit E or N gene sgRNA (***Figure 4H and Supplementary Figure 3J***). IHC staining of the lung tissues demonstrated high SARS-CoV-2 nucleocapsid protein expression in the control animals, whereas no viral antigen was detected in any of the scNP-immunized animals (***Figure 4I***). The sgRNA and histopathology data demonstrated that three doses of NTD-scNP or S2P-scNP immunization provided the same *in vivo* protection as RBD-scNP immunization, preventing SARS-CoV-2 infection in both lower and upper respiratory tracts.

### RBD-scNP, NTD-scNP and S2P-scNP as boosts for mRNA-LNP vaccine elicited various neutralizing antibody responses

We next assessed the efficacy of the RBD-scNP, NTD-scNP and S2P-scNP as boosts in macaques that received two priming doses of mRNA vaccine. Cynomolgus macaques (*n*=5) were immunized twice with 50 μg of S-2P-encoding, nucleoside-modified mRNA encapsulated in lipid nanoparticles (S-2P mRNA-LNP), which phenocopies the Pfizer/BioNTech and the Moderna COVID-19 vaccines.

Subsequently, macaques were boosted with RBD-, NTD- or S2P-scNPs (***Figure 4J***). Plasma antibody binding patterns were similar among the three groups until animals received the scNP boosting (***Supplementary Figure 4A***). Plasma antibodies targeting to ACE2-binding site and neutralizing epitopes were detected after the scNP boosting with cross-reactive antibodies in the DH1047 blocking assay being highest after RBD-scNP or S2P-NP boosting (***Supplementary Figure 4B-C***). BAL and nasal wash mucosal ACE2-blocking and DH1047-blocking activities tended to be low in magnitude in macaques primed with a Spike mRNA-LNP vaccine and boosted with RBD-scNP or S2P-scNP (***Supplementary Figure 4D-E***).

Serum neutralizing titers against the WA-1 strain pseudovirus were similar in the RBD-scNP-boosted group (GMT ID50 = 10,912.1) and S2P-scNP-boosted group (GMT ID50 = 7799.9) (***Figure 4K***), while the NTD-scNP-boosted group showed significantly lower titers (GMT ID50 = 3229.8; *p* = 0.027, exact Wilcoxon test). The same differences were also observed in other major variants (***Figure 4K***). In addition, in the RBD-scNP- and S2P-scNP-boosted groups, reduced ID50 titers were mostly seen for the Beta and Gamma variants, whereas in the NTD-scNP-boosted group, Alpha, Beta, Gamma, Delta, Iota and Kappa variants all showed >5-fold reduction of ID50 titers (***Figure 4L***).

Regarding Omicron neutralization, two doses of S2P mRNA-LNP immunization induced neutralizing antibodies to D614G with GMT ID50 of 7,814, which dropped 67-fold when testing for Omicron (GMT ID50=116) (***Figure 4M***). The RBD-scNP-boost at one month did not increase D614G neutralization titers, but raised Omicron neutralization titers to GMT ID50 of 374 (***Figure 4M***). Thus, NTD-scNP was an inferior boost of the S-2P mRNA-LNP vaccine compared to RBD- or S2P-scNPs, and the mRNA-LNP prime/RBD-scNP one-month boost showed reduced boosting capacity for neutralizing antibodies against Omicron, suggesting a longer boosting interval will be needed (Gagne et al., 2022).

### Protection of mRNA-LNP-primed and scNP-boosted macaques from SARS-CoV-2 challenge

Macaques that received mRNA-LNP prime and scNP boosts at one month post-mRNA-LNP primes were challenged with SARS-CoV-2 WA-1 strain after boosting. Four of five RBD-scNP-boosted monkeys and four of five of the S2P-scNP-boosted monkeys were completely protected from SARS-CoV-2 infection, showing no detectable E or N gene sgRNA in either BAL or nasal swab samples (***Figure 4N and Supplementary Figure 4F***). However, in the NTD-scNP boost group, N gene sgRNA was detected in BAL from three of five animals and in nasal swab samples from two of five animals (***Figure 4N***). Macaques that received mRNA-LNP prime and RBD-scNP boost had the lowest degree of lung inflammation (***Figure 4O***). In addition, no viral antigen was observed in lung tissues from either of the immunized groups as indicated by IHC staining for SARS-CoV-2 N protein (***Figure 4O***).

## DISCUSSION

In this study, the SARS-CoV-2 RBD has maintained conserved neutralizing epitopes among Beta, Delta and Omicron variants, despite the up to 15 RBD amino acid changes for the omicron variant relative to the WA-1 strain. Given the neutralization titers that have protected macaques in challenge studies, we speculate the RBD-scNP vaccine would protect against Omicron challenge with similar efficacy as shown here for the Beta variant. While SARS-CoV-2 continues to mutate during the ongoing pandemic, there are conserved RBD neutralizing epitopes among the SARS-CoV-2 variants. This result is supported by studies with a SARS-CoV-2 virus where 20 naturally occurring mutations were introduced into the spike protein, but the resultant virus was still sensitive to vaccine-induced polyclonal antibody responses (Schmidt et al., 2021). For present and future coverage of SARS-CoV-2 variants, it will be critical to induce a polyclonal response that targets conserved sites on the RBD. Monoclonal antibodies such as S2x259, S2K146 (Cameroni et al., 2021), DH1047 (Li *et al*., 2021a; Martinez *et al*., 2021a) and S309 (Pinto et al., 2020) have defined these key conserved sites upon which vaccines can be designed.

Adjuvants play essential roles in vaccine formulation to elicit strong protective immune responses (Coffman et al., 2010) and Alum is used in many currently approved vaccines (HogenEsch et al., 2018). Thus, it was encouraging to see that the RBD-NP vaccine was protective in NHPs when adsorbed to Alum. Compared to Alum, 3M-052-AF + Alum demonstrated superior capacities to elicit neutralizing antibodies against SARS-CoV-2 WA-1 live virus when formulated with SARS-CoV-2 RBD trimer in mice but not in rhesus macaques (Routhu et al., 2021). In addition, 3M-052-adjuvanted gp140 Env vaccine augmented neutralizing antibodies against tier 1A HIV-1 pseudovirus in rhesus macaques (Kasturi et al., 2020). 3M-052-AF + Alum is in clinical testing of HIV-1 vaccines (NCT04915768 and NCT04177355). Here we found that 3M-052-AF alone-adjuvanted RBD-scNPs induced not only superior systemic and mucosal antibody responses, but also higher titers of neutralizing antibodies than 3M-052- AF + Alum-adjuvanted vaccine, demonstrating that 3M-052-AF in the absence of Alum is an optimal adjuvant for scNP. One explanation for this difference could be antagonism between Th1-based immune pathways induced by 3M-052 (Smirnov et al., 2011) and Th2-based pathways induced by Alum (Marrack et al., 2009). Another potential explanation is that physicochemical considerations such as particle size or adsorption interactions between Alum and the RBD-scNP antigen and/or 3M-052-AF are impacting vaccine biodistribution, presentation, or cellular processing, thus affecting downstream immune responses. Such interactions are antigen-dependent (Fox et al., 2016), highlighting the importance of optimizing adjuvant formulation for each unique antigen (Fox et al., 2013; HogenEsch *et al*., 2018).

Coronavirus vaccines formulated with Alum have been reported to be associated with enhanced lung inflammation, particularly with killed vaccines (Arvin et al., 2020; Haynes et al., 2020). However, it is important to note that no enhancement of lung inflammation or virus replication was seen with RBD-scNP/Alum formulations. The RBD-scNP + 3M-052-AF group exhibited the highest neutralizing antibody titers and was the only group showing reduced severity of lung inflammation. That RBD-scNPs formulated with Alum alone protected monkeys after immunization raises the possibility of its use in pediatric populations.

While the RBD subunit has been shown to protect against SARS-CoV-2 challenge in animal models (Dai et al., 2020; Francica et al., 2021; Pino et al., 2021; Saunders *et al*., 2021; Yang et al., 2020), the NTD is also an immunodominant region for neutralizing antibodies (Cerutti et al., 2021; Chi et al., 2020; Li *et al*., 2021a; Martinez et al., 2021b; McCallum et al., 2021; Voysey et al., 2021). However, NTD is the site of multiple mutations and NTD antibody neutralization is, in general, less potent than RBD antibodies. Here, in this study, NTD-scNP-induced serum neutralizing antibodies were detected using a live SARS-CoV-2 virus but not using pseudovirus. The inconsistent neutralization activities of NTD antibodies in different neutralization assays have been previously observed (Chi *et al*., 2020; Li *et al*., 2021a). In addition, the NTD-scNP immunization likely induced not only neutralizing, but also non-neutralizing NTD antibodies (Li *et al*., 2021a). Serum from the NTD-scNP group did have ADCC activity, suggesting that non-neutralizing Fc receptor-mediated antibody activities could have been involved in protection. In this regard, we previously found that a non-neutralizing NTD antibody DH1052 provided partial protection after infusion into mice and non-human primates (Li *et al*., 2021a). Therefore, the complete protection conferred by scNP vaccination could be a result of both neutralizing and non-neutralizing Fc receptor-mediated antibody activities. Moreover, we found that boosting with RBD-scNP or S2P-scNP after S2P mRNA-LNPs priming afforded complete protection for monkeys after WA-1 challenge, while NTD-scNP boosting of S2P mRNA-LNPs priming led to incomplete protection. The mechanism of this latter finding is currently under investigation.

Our study has several limitations. First, our study did not evaluate the durability of vaccine-induced immune responses and protection against SARS-CoV-2 variants. Second, we did not set up longer time intervals between the second and the third booster vaccination, to mimic 4-6 month boosting interval in humans. Lastly, we challenged the animals with WA-1 strain, the Beta variant and the Delta variant; future *in vivo* protection studies will be required upon availability of viral stocks of other SARS-CoV-2 variants such as the Omicron variant.

Thus, our study demonstrates that scNP vaccines with SARS-CoV-2 spike or spike subunits confer potent protection in NHPs against WA-1, Beta and Delta variants, and that they induce neutralizing antibodies to all SARS-CoV-2 variants tested *in vitro*. These findings have important implications for virus escape from neutralizing antibody responses and for development of the next generation of COVID- 19 vaccines.

## MATERIALS AND METHODS

### Animals and immunizations

The study protocol and all veterinarian procedures were approved by the Bioqual IACUC per a memorandum of understanding with the Duke IACUC, and were performed based on standard operating procedures. Macaques studied were housed and maintained in an Association for Assessment and Accreditation of Laboratory Animal Care-accredited institution in accordance with the principles of the National Institutes of Health. All studies were carried out in strict accordance with the recommendations in the Guide for the Care and Use of Laboratory Animals of the National Institutes of Health in BIOQUAL (Rockville, MD). BIOQUAL is fully accredited by AAALAC and through OLAW, Assurance Number A-3086. All physical procedures associated with this work were done under anesthesia to minimize pain and distress in accordance with the recommendations of the Weatherall report, “The use of non-human primates in research.” Teklad 5038 Primate Diet was provided once daily by animal size and weight. The diet was supplemented with fresh fruit and vegetables. Fresh water was given ad libitum. All monkeys were maintained in accordance with the Guide for the Care and Use of Laboratory Animals.

Cynomolgus macaques were on average 8-9 years old and ranged from 2.75 to 8 kg in body weight. Male and female macaques per group were balanced when availability permitted. Studies were performed unblinded. The RBD-scNP, NTD-scNP and S2P-scNP immunogens were formulated with adjuvants as previously described (Fox *et al*., 2016) and given intramuscularly in the right and left quadriceps. In the first study, cynomolgus macaques (n=5) were immunized for three times with 100 μg of RBD-scNP, NTD-scNP and S2P-scNP adjuvanted with 5 μg of 3M-052 aqueous formulation admixed with 500 μg of alum in PBS. In the second study, cynomolgus macaques (n=5) were were immunized twice with 50 μg of S-2P mRNA-LNP (encoding the transmembrane spike protein stabilized with K986P and V987P mutations) and boosted once with 100 μg of RBD-scNP, NTD-scNP and S2P-scNP adjuvanted with 5 μg of 3M-052 aqueous formulation admixed with 500 μg of alum in PBS. In the third study, cynomolgus macaques were immunized for twice with 100 μg of RBD-scNP or recombinant soluble RBD with 5 μg of 3M-052 aqueous formulation admixed with 500 μg of alum in PBS. In the fourth study, macaques were divided into 8 groups (n=5 per group) as following: 1) control group: no immunization; 2) immunogen alone group: 100 μg of RBD-scNP; 3) RBD-scNP + 3M-052-Alum group: 100 μg of RBD-scNP + 5 μg of 3M-052 in aqueous formulation + 500 μg of Alum (i.e. aluminum ion); 4) 3M-052-Alum alone group: 5 μg of 3M-052 in aqueous formulation + 500 μg of Alum; 5) RBD-scNP + Alum group: 100 μg of RBD-scNP + 500 μg of Alum; 6) Alum alone group: 500 μg of Alum; 7) RBD-scNP + 3M-052-AF group: 100 μg of RBD-scNP + 5 μg of 3M-052 in aqueous formulation; 8) 3M-052-AF alone group: 5 μg of 3M-052 in aqueous formulation.

### SARS-CoV-2 viral challenge

For SARS-CoV-2 challenge, 10^5^ plaque-forming units (PFU) of SARS-CoV-2 virus Isolate USA- WA1/2020 (∼10^6^ TCID50) were diluted in 4 mL and were given by 1 mL intranasally and 3 mL intratracheally on Day 0. Biospecimens, including nasal swabs, BAL, plasma, and serum samples, were collected before immunization, after every immunization, before challenge, 2 days post-challenge and 4 days post-challenge. Animals were necropsied on Day 4 post-challenge, and lungs were collected for histopathology and immunohistochemistry (IHC) analysis.

### Recombinant protein production

The coronavirus ectodomain proteins were produced and purified as previously described (Li *et al*., 2021a; Saunders *et al*., 2021; Wrapp et al., 2020; Zhou et al., 2020). S-2P was stabilized by the introduction of 2 prolines at amino acid positions 986 and 987. Plasmids encoding SARS-CoV-2 and other coronavirus S-2P (Genscript) were transiently transfected in FreeStyle 293-F cells (Thermo Fisher) using Turbo293 (SpeedBiosystems) or 293Fectin (ThermoFisher). All cells were tested monthly for mycoplasma. The constructs contained an HRV 3C-cleavable C-terminal twinStrepTagII-8×His tag. On day 6, cell-free culture supernatant was generated by centrifugation of the culture and filtering through a 0.8-μm filter. Protein was purified from filtered cell culture supernatants by StrepTactin resin (IBA) and by size-exclusion chromatography using Superdex 200 (RBD and NTD) or Superose 6 (S-2P and ferritin) column (GE Healthcare) in 10 mM Tris pH=8, 500 mM NaCl. ACE2-Fc was expressed by transient transfection of Freestyle 293-F cells. ACE2-Fc was purified from cell culture supernatant by HiTrap protein A column chromatography and Superdex200 size-exclusion chromatography in 10 mM Tris pH8,150 mM NaCl. SARS-CoV-2 RBD and NTD were produced as previously described (Saunders *et al*., 2021; Zhou *et al*., 2020).

RBD-scNP, NTD-scNP, and S2P-scNP were produced by conjugating SARS-CoV-2 RBD to *H. pylori* ferritin nanoparticles using Sortase A as previously described (Saunders *et al*., 2021). Briefly, SARS-CoV-2 Wuhan strain RBD, NTD or S-2P (with a C-terminal foldon trimerization motif) was expressed with a sortase A donor sequence LPETGG encoded at its C terminus. C-terminal to the sortase A donor sequence was an HRV-3C cleavage site, 8×His tag and a twin StrepTagII (IBA). The proteins were expressed in Freestyle 293-F cells and purified by StrepTactin affinity chromatography and Superdex 200 or Superose 6 size-exclusion chromatography. *Helicobacter pylori* ferritin particles were expressed with a pentaglycine sortase A acceptor sequence encoded at its N terminus of each subunit. For affinity purification of ferritin particles, 6×His tags were appended C-terminal to a HRV3C cleavage site. Ferritin particles with a sortase A N-terminal tag were buffer exchanged into 50 mM Tris, 150 mM NaCl, 5 mM CaCl2, pH 7.5. Then 180 μM SARS-CoV-2 RBD was mixed with 120 μM of ferritin subunits and incubated with 100 μM of sortase A overnight at room temperature. Following incubation, conjugated particles were isolated from free ferritin or free RBD/NTD/S-2P by size-exclusion chromatography using a Superose 6 16/60 column.

### mRNA-LNP vaccine production

The S-2P mRNA was designed based on the SARS-CoV-2 spike (S) protein sequence (Wuhan-Hu-1) and encoded the full-length S with K986P and V987P amino acid substitutions. Production of the mRNA was performed as described earlier (Freyn et al., 2021; Freyn et al., 2020). Briefly, the codon-optimized S-2P gene was synthesized (Genscript) and cloned into an mRNA production plasmid. A T7-driven in vitro transcription reaction (Megascript, Ambion) using linearized plasmid template was performed to generate mRNA with 101 nucleotide long poly(A) tail. Capping of the mRNA was performed in concert with transcription through addition of a trinucleotide cap1 analog, CleanCap (TriLink) and m1Ψ-5’- triphosphate (TriLink) was incorporated into the reaction instead of UTP. Cellulose-based purification of S-2P mRNA was performed as described (Baiersdorfer et al., 2019). The S-2P mRNA was then tested on an agarose gel before storing at -20°C. The cellulose-purified m1Ψ-containing S-2P mRNA was encapsulated in LNPs using a self-assembly process as previously described wherein an ethanolic lipid mixture of ionizable cationic lipid, phosphatidylcholine, cholesterol and polyethylene glycol-lipid was rapidly mixed with an aqueous solution containing mRNA at acidic pH (Maier et al., 2013). The RNA-loaded particles were characterized and subsequently stored at 80 °C at a concentration of 1 mg/ml.

### Antibody Binding ELISA

For binding ELISA, 384-well ELISA plates were coated with 2 μg/mL of antigens in 0.1 M sodium bicarbonate overnight at 4°C. Plates were washed with PBS + 0.05% Tween 20 and blocked with blocked with assay diluent (PBS containing 4% (w/v) whey protein, 15% Normal Goat Serum, 0.5% Tween-20, and 0.05% Sodium Azide) at room temperature for 1 hour. Plasma or mucosal fluid were serially diluted threefold in superblock starting at a 1:30 dilution. Nasal fluid was started from neat, whereas BAL fluid was concentrated ten-fold. To concentrate BAL, individual BAL aliquots from the same macaque and same time point were pooled in 3-kDa MWCO ultrafiltration tubes (Sartorious, catalog # VS2091).

Pooled BAL was concentrated by centrifugation at 3,500 rpm for 30 min or until volume was reduced by a factor of 10. The pool was then aliquoted and frozen at −80 °C until its use in an assay. Serially diluted samples were added and incubated for 1 hour, followed by washing with PBS-0.1% Tween 20. HRP-conjugated goat anti-human IgG secondary Ab (SouthernBiotech, catalog# 2040-05) was diluted to 1:10,000 and incubated at room temperature for 1 hour. These plates were washed four times and developed with tetramethylbenzidine substrate (SureBlue Reserve- KPL). The reaction was stopped with 1 M HCl, and optical density at 450 nm (OD450) was determined.

### ACE2 and neutralizing antibody blocking assay

ELISA plates were coated as stated above with 2 μg/mL recombinant ACE-2 protein or neutralizing antibodies, then washed and blocked with 3% BSA in 1x PBS. While assay plates blocked, plasma or mucosal samples were diluted as stated above, only in 1% BSA with 0.05% Tween-20. In a separate dilution plate spike-2P protein was mixed with the antibodies at a final concentration equal to the EC50 at which spike binds to ACE-2 protein. The mixture was incubated at room temperature for 1 hour. Blocked assay plates were then washed and the antibody-spike mixture was added to the assay plates for a period of 1 hour at room temperature. Plates were washed and a polyclonal rabbit serum against the same spike protein (nCoV-1 nCoV-2P.293F) was added for 1 hour, washed and detected with goat anti rabbit-HRP (Abcam catalog # ab97080) followed by TMB substrate. The extent to which antibodies were able to block the binding spike protein to ACE-2 or neutralizing antibodies was determined by comparing the OD of antibody samples at 450 nm to the OD of samples containing spike protein only with no antibody. The following formula was used to calculate percent blocking: blocking % = (100 - (OD sample/OD of spike only)*100).

### Pseudotyped SARS-CoV-2 neutralization assay

Neutralization of SARS-CoV-2 Spike-pseudotyped virus was performed by adopting an infection assay described previously (Korber et al., 2020) with lentiviral vectors and infection in 293T/ACE2.MF (the cell line was kindly provided by Drs. Mike Farzan and Huihui Mu at Scripps). Cells were maintained in DMEM containing 10% FBS and 50 µg/ml gentamicin. An expression plasmid encoding codon- optimized full-length spike of the Wuhan-1 strain (VRC7480), was provided by Drs. Barney Graham and Kizzmekia Corbett at the Vaccine Research Center, National Institutes of Health (USA). Mutations were introduced into VRC7480 either by site-directed mutagenesis using the QuikChange Lightning Site- Directed Mutagenesis Kit from Agilent Technologies (Catalog # 210518), or were created by spike gene synthesized by GenScript using the spike sequence in VRC7480 as template. All mutations were confirmed by full-length spike gene sequencing by Sanger Sequencing, using Sequencher and SnapGene for sequence analyses. D614G spike mutation: D614G; Alpha (B.1.1.7) spike mutations: Δ69-70, Δ144, N501Y, A570D, D614G, P681H, T716I, S982A, D1118H; Beta (B.1.351) spike mutations: L18F, D80A, D215G, Δ242-244, R246I, K417N, E484K, N501Y, D614G, A701V; Delta (B.1.617 AY.3) spike mutations: T19R, G142D, Δ156-157, R158G, L452R, T478K, D614G, P681R, D950N; Epsilon (B.1.429) spike mutations: S13I, W152C, L452R, D614G; Omicron (B.1.1.529) spike mutations: A67V, Δ69-70, T95I, G142D, Δ143-145, Δ211, L212I, +214EPE, G339D, S371L, S373P, S375F, K417N, N440K, G446S, S477N, T478K, E484A, Q493R, G496S, Q498R, N501Y, Y505H, T547K, D614G, H655Y, N679K, P681H, N764K, D796Y, N856K, Q954H, N969K, L981F. Pseudovirions were produced in HEK 293T/17 cells (ATCC cat. no. CRL-11268) by transfection using Fugene 6 (Promega, Catalog #E2692). Pseudovirions for 293T/ACE2 infection were produced by co-transfection with a lentiviral backbone (pCMV ΔR8.2) and firefly luciferase reporter gene (pHR’ CMV Luc) (Naldini et al., 1996). Culture supernatants from transfections were clarified of cells by low-speed centrifugation and filtration (0.45 µm filter) and stored in 1 ml aliquots at -80 °C. A pre-titrated dose of virus was incubated with 8 serial 3-fold or 5-fold dilutions of mAbs in duplicate in a total volume of 150 µl for 1 hr at 37 °C in 96-well flat- bottom poly-L-lysine-coated culture plates (Corning Biocoat). Cells were suspended using TrypLE express enzyme solution (Thermo Fisher Scientific) and immediately added to all wells (10,000 cells in 100 µL of growth medium per well). One set of 8 control wells received cells + virus (virus control) and another set of 8 wells received cells only (background control). After 66-72 hrs of incubation, medium was removed by gentle aspiration and 30 µL of Promega 1x lysis buffer was added to all wells. After a 10-minute incubation at room temperature, 100 µl of Bright-Glo luciferase reagent was added to all wells. After 1-2 minutes, 110 µl of the cell lysate was transferred to a black/white plate (Perkin-Elmer).

Luminescence was measured using a PerkinElmer Life Sciences, Model Victor2 luminometer. Neutralization titers are the mAb concentration (IC50/IC80) at which relative luminescence units (RLU) were reduced by 50% and 80% compared to virus control wells after subtraction of background RLUs. Negative neutralization values are indicative of infection-enhancement. Maximum percent inhibition (MPI) is the reduction in RLU at the highest mAb concentration tested.

Another protocol was used to test plasma neutralization against pseudoviruses of SARS-CoV-2 WA- 1 strain and variants. Human codon-optimized cDNA encoding SARS-CoV-2 spike glycoproteins of various strains were synthesized by GenScript and cloned into eukaryotic cell expression vector pcDNA 3.1 between the BamHI and XhoI sites. Pseudovirions were produced by co-transfection of Lenti-X 293T cells with psPAX2(gag/pol), pTrip-luc lentiviral vector and pcDNA 3.1 SARS-CoV-2-spike-deltaC19, using Lipofectamine 3000. The supernatants were collected at 48 h after transfection and filtered through 0.45-μm membranes and titrated using HEK293T cells that express ACE2 and TMPRSS2 protein (293T- ACE2-TMPRSS2 cells). For the neutralization assay, 50 μl of SARS-CoV-2 spike pseudovirions were pre-incubated with an equal volume of medium containing serum at varying dilutions at room temperature for 1 h, then virus-antibody mixtures were added to 293T-ACE2-TMPRSS2 cells in a 96-well plate. After a 3-h incubation, the inoculum was replaced with fresh medium. Cells were lysed 24 h later, and luciferase activity was measured using luciferin. Controls included cell-only control, virus without any antibody control and positive control sera. Neutralization titres are the serum dilution (ID50 or ID80) at which relative luminescence units (RLU) were reduced by 50% or 80%, respectively, compared to virus control wells after subtraction of background RLUs.

### Live SARS-CoV-2 neutralization assays

The SARS-CoV-2 virus (Isolate USA-WA1/2020, NR-52281) was deposited by the Centers for Disease Control and Prevention and obtained through BEI Resources, NIAID, NIH. SARS-CoV-2 Micro- neutralization (MN) assays were adapted from a previous study (Berry et al., 2004). In short, sera or purified Abs are diluted two-fold and incubated with 100 TCID50 virus for 1 hour. These dilutions are used as the input material for a TCID50. Each batch of MN includes a known neutralizing control Ab (Clone D001; SINO, CAT# 40150-D001). Data are reported as the concentration at which 50% of input virus is neutralized. A known neutralizing control antibody is included in each batch run (Clone D001; SINO, CAT# 40150-D001). GraphPad Prism was used to determine ID50 values.

### Spike protein-expressing cell antibody binding assay

The cell antibody binding assay was performed as previously described (Pino et al., 2021). Briefly, target cells were derived by transfection with plasmids designed to express the SARS-CoV-2 D614 Spike protein with a c-terminus flag tag (kindly provided by Dr. Farzan, Addgene plasmid no. 156420 (Zhang et al., 2020)). Cells not transfected with any plasmid (mock transfected) were used as a negative control condition. After resuspension, washing and counting, 1x10^5^ Spike-transfected target cells were dispensed into 96-well V-bottom plates and incubated with six serial dilutions of macaque plasma starting at 1:50 dilution. Mock transfected cells were used as a negative infection control. After 30 minutes incubation at 37°C, cells are washed twice with 250 μL/well of PBS, stained with vital dye (Live/Dead Far Red Dead Cell Stain, Invitrogen) to exclude nonviable cells from subsequent analysis, washed with Wash Buffer (1%FBS-PBS; WB), permeabilized with CytoFix/CytoPerm (BD Biosciences), and stained with 1.25 µg/mL anti-human IgG Fc-PE/Cy7 (Clone HP6017; Biolegend) and 5 µg/mL anti-flag-FITC (clone M2; Sigma Aldrich) in the dark for 20 minutes at room temperature. After three washes with Perm Wash (BD Biosciences), the cells were resuspended in 125 μL PBS-1% paraformaldehyde. Samples were acquired within 24 h using a BD Fortessa cytometer and a High Throughput Sampler (HTS, BD Biosciences). Data analysis was performed using FlowJo 10 software (BD Biosciences). A minimum of 50,000 total events were acquired for each analysis. Gates were set to include singlet, live, flag+ and IgG+ events. All final data represent specific binding, determined by subtraction of non-specific binding observed in assays performed with mock-transfected cells.

### Antibody-dependent NK cell degranulation assay

Cell-surface expression of CD107a was used as a marker for NK cell degranulation, a prerequisite process for ADCC (Ferrari et al., 2011), was performed as previously described (Pino et al., 2021).

Briefly, target cells were either Vero E6 cells after a 2 day-infection with SARS-CoV-2 USA-WA1/2020 or 293T cells 2-days post transfection with a SARS-CoV-2 S protein (D614) expression plasmid. NK cells were purified from peripheral blood of a healthy human volunteer by negative selection (Miltenyi Biotech), and were incubated with target cells at a 1:1 ratio in the presence of diluted plasma or monoclonal antibodies, Brefeldin A (GolgiPlug, 1 μl/ml, BD Biosciences), monensin (GolgiStop, 4μl/6mL, BD Biosciences), and anti-CD107a-FITC (BD Biosciences, clone H4A3) in 96-well flat bottom plates for 6 hours at 37°C in a humidified 5% CO2 incubator. NK cells were then recovered and stained for viability prior to staining with CD56-PECy7 (BD Biosciences, clone NCAM16.2), CD16-PacBlue (BD Biosciences, clone 3G8), and CD69-BV785 (Biolegend, Clone FN50). Flow cytometry data analysis was performed using FlowJo software (v10.8.0). Data is reported as the % of CD107A+ live NK cells (gates included singlets, lymphocytes, aqua blue-, CD56+ and/or CD16+, CD107A+). All final data represent specific activity, determined by subtraction of non-specific activity observed in assays performed with mock-infected cells and in absence of antibodies.

### Viral RNA Extraction and Subgenomic mRNA quantification

SARS-CoV-2 E gene and N gene subgenomic mRNA (sgRNA) was measured by a one-step RT-qPCR adapted from previously described methods (Wolfel et al., 2020; Yu et al., 2020). To generate standard curves, a SARS-CoV-2 E gene sgRNA sequence, including the 5’UTR leader sequence, transcriptional regulatory sequence (TRS), and the first 228 bp of E gene, was cloned into a pcDNA3.1 plasmid. For generating SARS-CoV-2 N gene sgRNA, the E gene was replaced with the first 227 bp of N gene. The recombinant pcDNA3.1 plasmid was linearized, transcribed using MEGAscript T7 Transcription Kit (ThermoFisher, catalog # AM1334), and purified with MEGAclear Transcription Clean-Up Kit (ThermoFisher, catalog # AM1908). The purified RNA products were quantified on Nanodrop, serial diluted, and aliquoted as E sgRNA or N sgRNA standards.

A QIAsymphony SP (Qiagen, Hilden, Germany) automated sample preparation platform along with a virus/pathogen DSP midi kit. RNA extracted from animal samples or standards were then measured in Taqman custom gene expression assays (ThermoFisher). For these assays we used TaqMan Fast Virus 1- Step Master Mix (ThermoFisher, catalog # 4444432) and custom primers/probes targeting the E gene sgRNA (forward primer: 5’ CGA TCT CTT GTA GAT CTG TTC TCE 3’; reverse primer: 5’ ATA TTG CAG CAG TAC GCA CAC A 3’; probe: 5’ FAM-ACA CTA GCC ATC CTT ACT GCG CTT CG-BHQ1 3’) or the N gene sgRNA (forward primer: 5’ CGA TCT CTT GTA GAT CTG TTC TC 3’; reverse primer: 5’ GGT GAA CCA AGA CGC AGT AT 3’; probe: 5’ FAM-TAA CCA GAA TGG AGA ACG CAG TGG G-BHQ1 3’). RT-qPCR reactions were carried out on CFX384 Touch Real-Time PCR System (Bio-Rad) using a program below: reverse transcription at 50°C for 5 minutes, initial denaturation at 95°C for 20 seconds, then 40 cycles of denaturation-annealing-extension at 95°C for 15 seconds and 60°C for 30 seconds. Standard curves were used to calculate E or N sgRNA in copies per ml; the limit of detections (LOD) for both E and N sgRNA assays were 12.5 copies per reaction or 150 copies per mL of BAL/nasal swab.

### Histopathology

Lung specimen from nonhuman primates were fixed in 10% neutral buffered formalin, processed, and blocked in paraffin for histological analysis. All samples were sectioned at 5 µm and stained with hematoxylin-eosin (H&E) for routine histopathology. Sections were examined under light microscopy using an Olympus BX51 microscope and photographs were taken using an Olympus DP73 camera.

Samples were scored by a board-certified veterinary pathologist in a blinded manner. The representative images are to characterize the types and arrangement of inflammatory cells, while the scores show the relative severity of the tissue section.

### Immunohistochemistry (IHC)

Staining for SARS-CoV-2 antigen was achieved on the Bond RX automated system with the Polymer Define Detection System (Leica) used per manufacturer’s protocol. Tissue sections were dewaxed with Bond Dewaxing Solution (Leica) at 72°C for 30 min then subsequently rehydrated with graded alcohol washes and 1x Immuno Wash (StatLab). Heat-induced epitope retrieval (HIER) was performed using Epitope Retrieval Solution 1 (Leica), heated to 100°C for 20 minutes. A peroxide block (Leica) was applied for 5 min to quench endogenous peroxidase activity prior to applying the SARS-CoV-2 antibody (1:2000, GeneTex, GTX135357). Antibodies were diluted in Background Reducing Antibody Diluent (Agilent). The tissue was subsequently incubated with an anti-rabbit HRP polymer (Leica) and colorized with 3,3’-Diaminobenzidine (DAB) chromogen for 10 min. Slides were counterstained with hematoxylin.

### Negative-stain electron microscopy

Samples diluted to 200 µg/ml with 5 g/dl Glycerol in HBS (20 mM HEPES, 150 mM NaCl pH 7.4) buffer containing 8 mM glutaraldehyde. After 5 min incubation, glutaraldehyde was quenched by adding sufficient 1 M Tris stock, pH 7.4, to give 75 mM final Tris concentration and incubated for 5 min. Quenched sample was applied to a glow-discharged carbon-coated EM grid for 10-12 second, then blotted, and stained with 2 g/dL uranyl formate for 1 min, blotted and air-dried. Grids were examined on a Philips EM420 electron microscope operating at 120 kV and nominal magnification of 82,000x, and images were collected on a 4 Mpix CCD camera at 4 Å/pixel. Images were analyzed by 2D class averages using standard protocols with Relion 3.0 (Zivanov et al., 2018). The Relion reference needs to be added to the references. Also let us know which lot# you are shown in this manuscript so we can double check whether images are collected with old or new camera. Since the magnification and Å/pixel is different.

### Mouse immunization and challenge

Eleven-month-old female BALB/c mice were purchased from Envigo (#047) and were used for the SARS-CoV, SARS-CoV-2 WA-1, SARS-CoV-2 B.1.351, and RsSHC014-CoV protection experiments. The study was carried out in accordance with the recommendations for care and use of animals by the Office of Laboratory Animal Welfare (OLAW), National Institutes of Health and the Institutional Animal Care and Use Committee (IACUC) of University of North Carolina (UNC permit no. A-3410-01).

Animals were housed in groups of five and fed standard chow diets. Virus inoculations were performed under anesthesia and all efforts were made to minimize animal suffering. Mice were intramuscularly immunized with RBD-scNP formulated with 3M-052-AF + Alum or GLA-SE. For the SARS-CoV-2 WA-1 and RsSHC014 study, mice were immunized on week 0 and 2, and challenged on week 7. For the SARS-CoV-2 B.1.351 and SARS-CoV-1 study, mice were immunized on week 0 and 4, and challenged on week 6. All mice were anesthetized and infected intranasally with 1 × 10^4^ PFU/ml of SARS-CoV MA15, 1 × 10^4^ PFU/ml of SARS-CoV-2 WA1- MA10 or B.1.351-MA10, 1 × 10^4^ PFU/ml RsSHC014, which have been described previously (Leist et al., 2020; Martinez *et al*., 2021a; Martinez *et al*., 2021b; Menachery et al., 2015; Roberts et al., 2007). Mice were weighted daily and monitored for signs of clinical disease, and selected groups were subjected to daily whole-body plethysmography. For all mouse studies, groups of n=10 mice were included per arm of the study. Lung viral titers and weight loss were measured from individual mice per group.

### Biocontainment and biosafety

Studies were approved by the UNC Institutional Biosafety Committee approved by animal and experimental protocols in the Baric laboratory. All work described here was performed with approved standard operating procedures for SARS-CoV-2 in a biosafety level 3 (BSL-3) facility conforming to requirements recommended in the Microbiological and Biomedical Laboratories, by the U.S. Department of Health and Human Service, the U.S. Public Health Service, and the U.S. Center for Disease Control and Prevention (CDC), and the National Institutes of Health (NIH).

### Statistics Analysis

Data were plotted using Prism GraphPad 8.0. Wilcoxon rank sum exact test was performed to compare differences between groups with p-value < 0.05 considered significant using SAS 9.4 (SAS Institute, Cary, NC). The Benjamini-Hochberg correction (Benjamini and Hochberg, 1995) was used to adjust the *p*-values for multiple comparisons.

## DATA AVAILABILITY

The data that support the findings of this study are available from the corresponding authors upon reasonable request.

## AUTHOR CONTRIBUTIONS

D.L. performed RNA assays, analyzed the data and wrote the manuscript; D.R.M., A.S., and R.S.B. performed or supervised the mouse challenge studies; L.L.S., M.B., W.E., A.N. performed monkey studies and managed monkey samples; H.C., E.L., A.B. performed protein production; M.B., R.P. carried out ELISA assays; T.H.O., G.D.S., D.C.M., A.E., H.G. and S.K. carried out neutralization assays; D.L., C.T.M., T.N.D., M.G., D.C.D. designed or performed subgenomic RNA assays; K.W.B., M.M., B.M.N., I.N.M. performed histopathology analysis; D.M., G.F., D.W.C., and S.S. performed staining and ADCC assays; K.M. and R.J.E. performed negative staining EM; R.W.R. and Y.W. performed statistical analyses; M.A.T. selected and provided adjuvant; C.B.F. formulated 3M-052 in alum; N.P. and D.W. provided the mRNA-LNP vaccine; M.A.T. selected and provided adjuvant; C.B.F. formulated 3M-052; M.G.L., H.A. and R.S. evaluated and supervised monkey studies; K.O.S. designed the antigens, supervised the protein production, and edited the manuscript; B.F.H. designed and managed the study, reviewed all data and wrote and edited the manuscript. All authors edited and approved the manuscript.

## Supporting information

Supplementary Raw Data for Histopathologic Analysis

## ACKNOWLEDGEMENTS

We thank Margaret Deyton, Victoria Gee-Lai, Aja Sanzone, Nolan Jamieson, Lena Smith, Nicole De Naeyer and Conor Anderson for technical assistance. We thank Elizabeth Donahue, Cynthia Nagle and Kelly Soderberg for program management. We thank John Harrison, Alex Granados, Adrienne Goode, Anthony Cook, Alan Dodson, Katelyn Steingrebe, Bridget Bart, Laurent Pessaint, Alex VanRy, Daniel Valentin, Amanda Strasbaugh, and Mehtap Cabus for assistance with macaque studies. This work was supported by funds from the State of North Carolina with funds from the federal CARES Act; NIH, NIAID, DAIDS grant AI142596 (B.F.H.), AI158571 (B.F.H.), UC6-AI058607, G20-AI167200 (G.D.S.); the Ting Tsung & Wei Fong Chao Foundation (B.F.H.); Hanna H Gray Fellowship from the Howard Hughes Medical Institute and a Postdoctoral Enrichment Award from the Burroughs Wellcome Fund (D.R.M.).

---

**Supplementary Figure 1.**
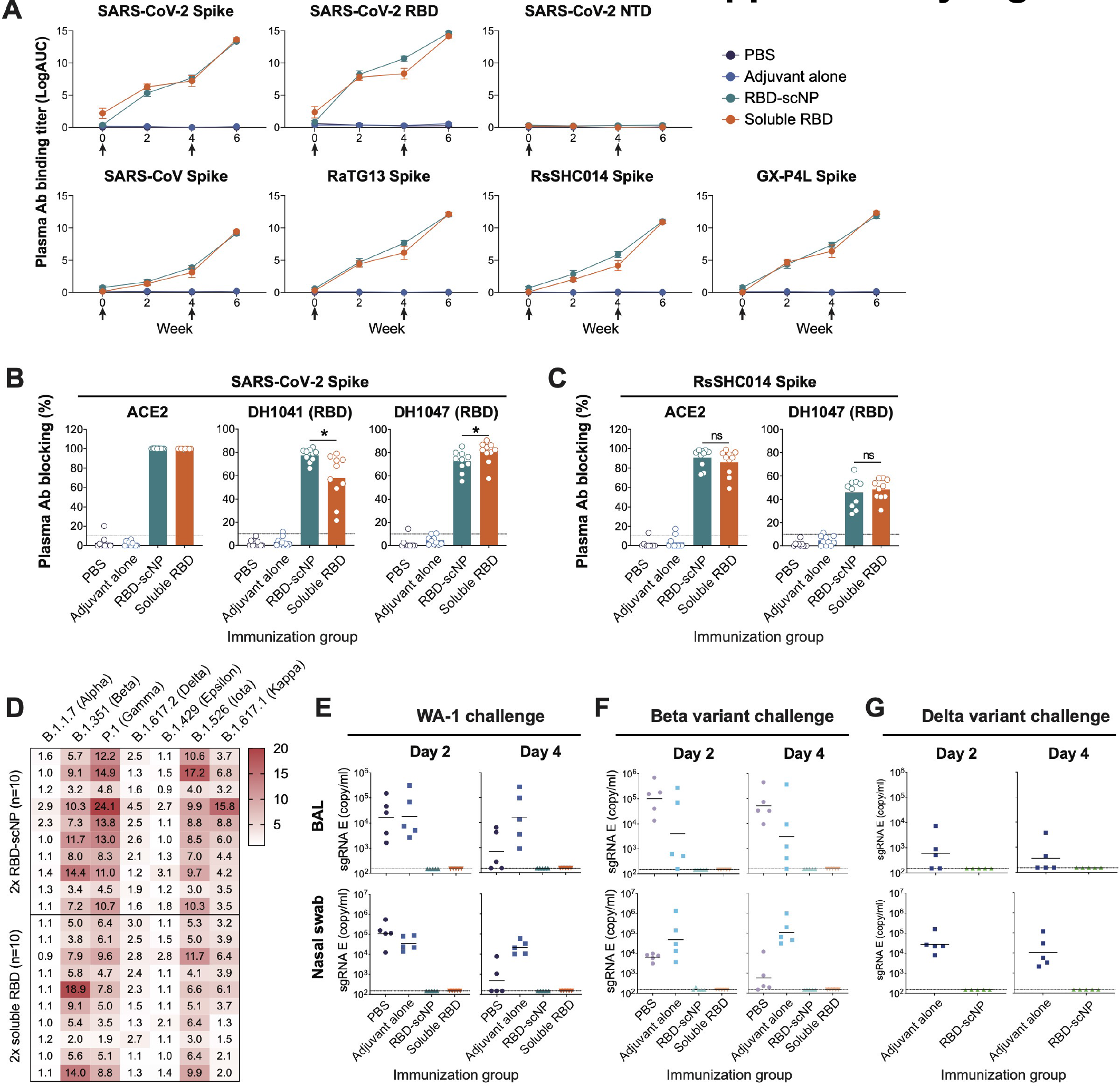
RBD-scNP elicited higher titers of neutralizing antibodies than soluble RBD. Related to Figure 2. **(A)** Plasma antibody binding titers to SARS-CoV-2 spike, RBD and NTD, as well as recombinant spike proteins of SARS-CoV, bat CoVs RaTG13, RsSHC014, and pangolin CoV GX-P4L. ELISA binding titers are shown as mean ± SEM of log area-under-curve (AUC). **(B-C)** Plasma antibody (post-2^nd^ immunization) blocking activity. ELISA was performed to test plasma antibodies blocking ACE2, human RBD neutralizing antibodies DH1041 and DH1047 binding to SARS-CoV-2 spike protein (B), or blocking ACE2 and DH1047 binding to RsSHC014 spike protein (C). Data are expressed as % blocking of ACE or the indicated antibody by 1:50 diluted plasma samples. **(D)** Fold reduction of plasma antibody ID_50_ titers against pseudoviruses of SARS-CoV-2 variants in 293T-ACE2-TMPRSS2 cells, compared to the titers against WA-1. **(E-G)** SARS-CoV-2 E gene sgRNA in BAL and nasal swab samples from the WA-1 (E), Beta variant (F), and Delta variant (G) challenged monkeys. Dashed line indicates limit of the detection. ns, not significant, *P<0.05, Wilcoxon rank sum exact test.

**Supplementary Figure 2.**
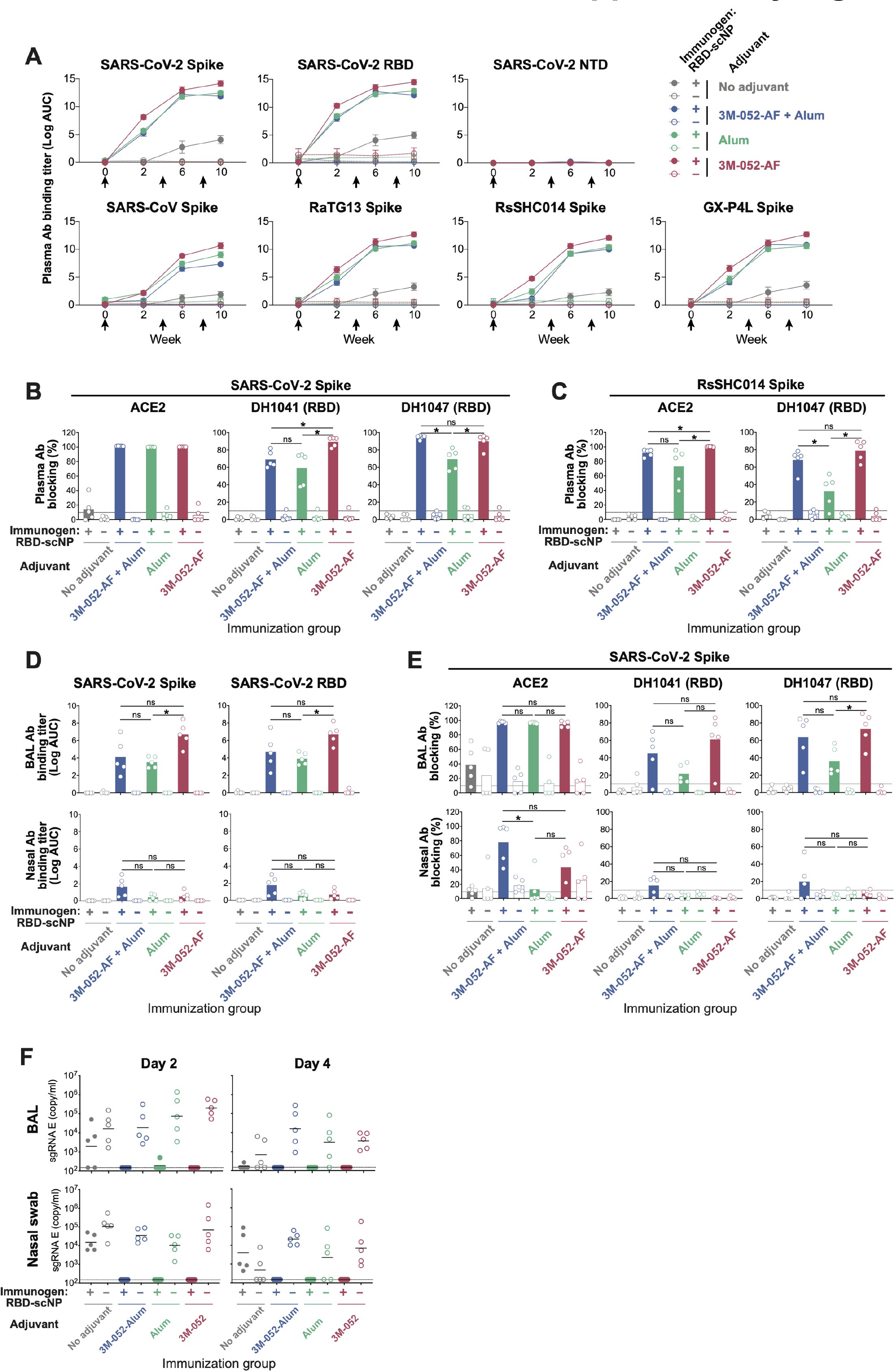
Serum and mucosal antibody responses elicited by RBD-scNP formulated with three different adjuvants. Related to Figure 3. **(A)** Plasma antibody binding titers to SARS-CoV-2 spike, RBD and NTD, as well as recombinant spike proteins of SARS-CoV, bat CoV RaTG13, RsSHC014, and pangolin CoV GX-P4L. ELISA binding titers are shown as mean ± SEM of log area-under-curve (AUC). **(B-C)** Plasma antibody (post-2^nd^ immunization) blocking activity. ELISA was performed to test plasma antibodies blocking ACE2, human RBD neutralizing antibodies DH1041 and DH1047 binding to SARS-CoV-2 spike protein (B), or blocking ACE2 and DH1047 binding to RsSHC014 spike protein (C). Data are expressed as % blocking of ACE or the indicated antibody by 1:50 diluted plasma samples. **(D-E)** Mucosal antibody binding and blocking activities after the 3^rd^ immunization. ELISA was performed to test 10x concentrated BAL or unconcentrated nasal wash samples binding to SARS- CoV-2 spike and RBD (D), or blocking ACE2, DH1041 and DH1047 binding to SARS-CoV-2 spike protein (E). Binding titers are expressed as log AUC, and blocking activities are shown as %blocking of ACE or the indicated antibody. **(F)** SARS-CoV-2 E gene sgRNA in BAL and nasal swab samples from the WA-1 challenged monkeys. Dashed line indicates limit of the detection. ns, not significant, *P<0.05, Wilcoxon rank sum exact test.

**Supplementary Figure 3.**
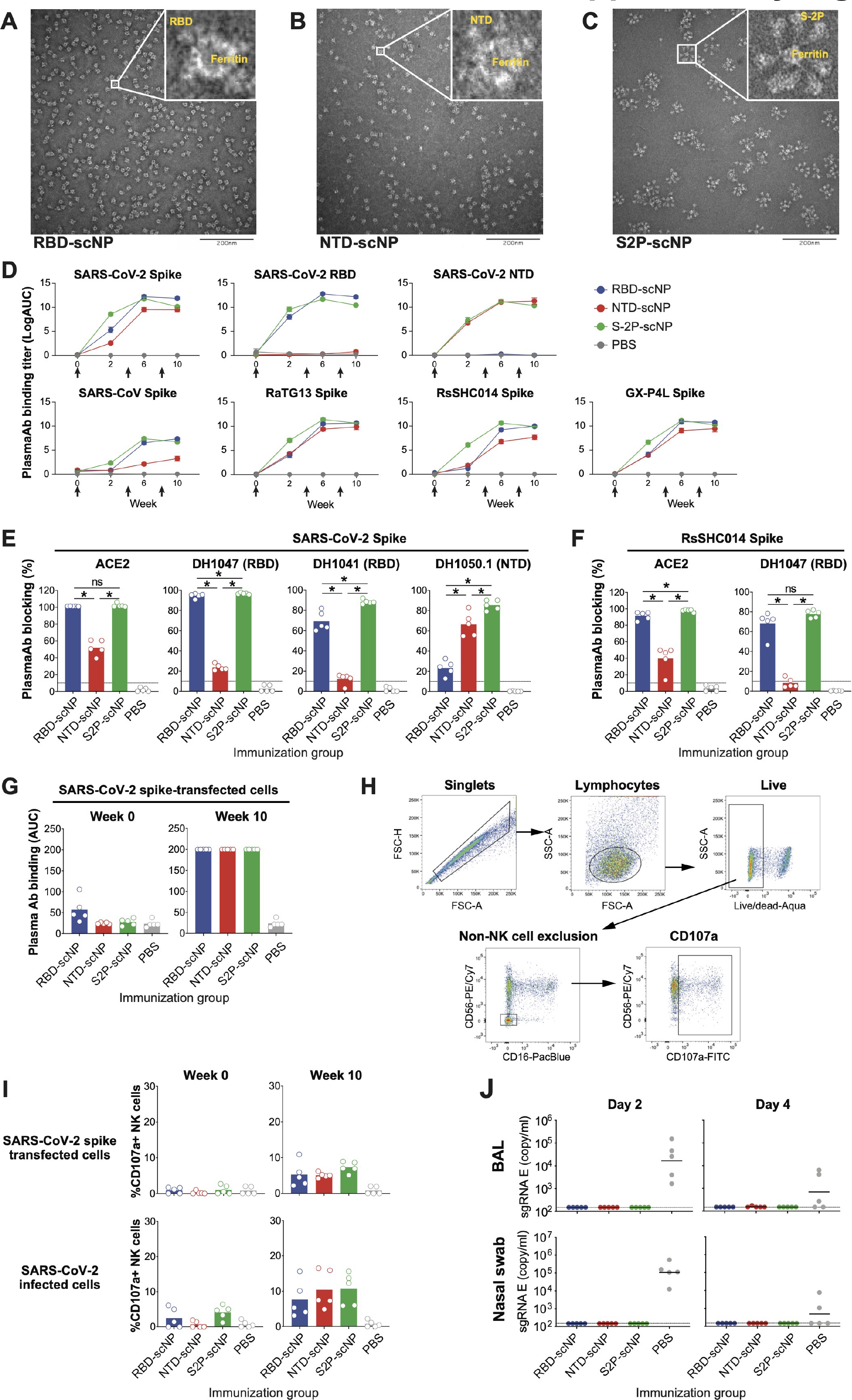
RBD-scNP, NTD-scNP and S2P-scNP induced spike-binding antibodies and mediated antibody-dependent cellular cytotoxicity (ADCC). Related to Figure 4. **(A-C)** Negative stain electron microscopy imaging of RBD-scNP (A), NTD-scNP (B), and S2P- scNP (C). The inset shows zoomed-in image of representative scNPs. **(D)** Plasma antibody binding titers to SARS-CoV-2 spike, RBD and NTD, as well as recombinant spike proteins of SARS-CoV, bat CoV RaTG13, RsSHC014, and pangolin CoV GX-P4L. ELISA binding titers are shown as mean ± SEM of log area-under-curve (AUC). **(E-F)** Plasma antibody (post-2^nd^ immunization) blocking activity. ELISA was performed to test plasma antibodies blocking ACE2, human RBD neutralizing antibodies DH1041 and DH1047, human NTD neutralizing antibodies DH1050.1 binding to SARS-CoV-2 spike protein (E), or blocking ACE2 and DH1047 binding to RsSHC014 spike protein (F). Data are expressed as % blocking of ACE or the indicated antibody by 1:50 diluted plasma samples. **(G)** Pre-immunization (week 0) and pre-challenge (week 10, post-3rd immunization) plasma antibodies binding on SARS-CoV-2 spike-transfected 293T cells tested by cell surface staining. **(H)** The gating strategy for the NK cell degranulation ADCC assay. Purified human NK cells were mixed with SARS-CoV-2 spike-transfected cells or SARS-CoV-2 infected cells in the presence of 1:50 diluted plasma samples. NK cell degranulation was detected based on CD107a expression. **(I)** RBD-scNP-, NTD-scNP- and S2P-scNP-induced antibodies mediated ADCC. The percentages of CD107a+ NK cells were shown when NK cells were assayed with plasma antibodies (week 0 and week 10) in SARS-CoV-2 spike transfected 293T cells (top row) or SARS-CoV-2 infected Vero E6 cells cells (botton row). **(J)** SARS-CoV-2 E gene sgRNA in BAL and nasal swab samples from WA-1 challenged monkeys. Dashed line indicates limit of the detection.

**Supplementary Figure 4.**
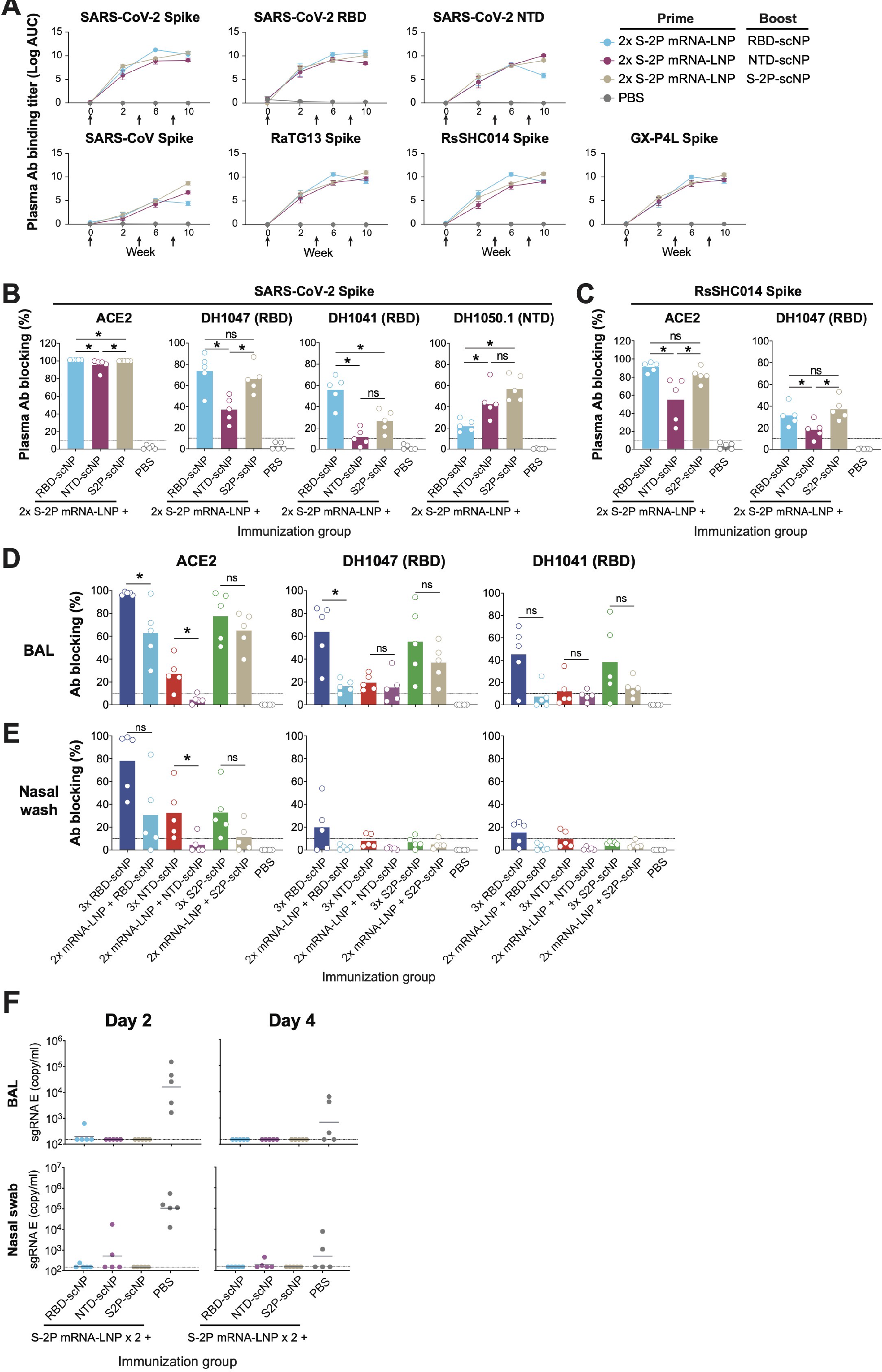
Antibody responses elicited by scNP vaccines as a booster vaccination in macaques that received two doses of S-2P mRNA-LNP vaccine. Related to Figure 4. **(A)** Plasma antibody binding titers to SARS-CoV-2 spike, RBD and NTD, as well as recombinant spike proteins of SARS-CoV, bat CoV RaTG13, RsSHC014, and pangolin CoV GX-P4L. ELISA binding titers are shown as mean ± SEM of log area-under-curve (AUC). **(B-C)** Plasma antibody (post-2^nd^ immunization) blocking activity. ELISA was performed to test plasma antibodies blocking ACE2, human RBD neutralizing antibodies DH1041 and DH1047, human NTD neutralizing antibodies DH1050.1 binding to SARS-CoV-2 spike protein (B), or blocking ACE2 and DH1047 binding to RsSHC014 spike protein (C). Data are expressed as % blocking of ACE or the indicated antibody by 1:50 diluted plasma samples. **(D-E)** Comparison of mucosal antibody blocking activities induced by 3 doses of scNP vaccination or 2 doses of S2P mRNA-LNP + 1 dose of scNP vaccination. ELISA for 10x concentrated BAL samples (D) and neat nasal wash samples (E) blocking the binding of ACE2 or neutralizing antibody (DH1041 or DH1047) on SARS-CoV-2 spike were performed. Data are expressed as % blocking of ACE or the indicated antibody by mucosal samples. **(F)** SARS-CoV-2 E gene sgRNA in BAL and nasal swab samples from WA-1 challenged monkeys. Dashed line indicates limit of the detection. ns, not significant, *P<0.05, Wilcoxon rank sum exact test.

## REFERENCES

1. Arvin, A.M., Fink, K., Schmid, M.A., Cathcart, A., Spreafico, R., Havenar-Daughton, C., Lanzavecchia, A., Corti, D., and Virgin, H.W. (2020). A perspective on potential antibody-dependent enhancement of SARS-CoV-2. Nature 584, 353–363. 10.1038/s41586-020-2538-8.

2. Baden, L.R., El Sahly, H.M., Essink, B., Kotloff, K., Frey, S., Novak, R., Diemert, D., Spector, S.A., Rouphael, N., Creech, C.B., et al. (2021). Efficacy and Safety of the mRNA-1273 SARS-CoV-2 Vaccine. N Engl J Med 384, 403–416. 10.1056/NEJMoa2035389.

3. Baiersdorfer, M., Boros, G., Muramatsu, H., Mahiny, A., Vlatkovic, I., Sahin, U., and Kariko, K. (2019). A Facile Method for the Removal of dsRNA Contaminant from In Vitro-Transcribed mRNA. Mol Ther Nucleic Acids 15, 26–35. 10.1016/j.omtn.2019.02.018.

4. Benjamini, Y., and Hochberg, Y. (1995). Controlling the False Discovery Rate: A Practical and Powerful Approach to Multiple Testing. Journal of the Royal Statistical Society. Series B (Methodological) 57, 289–300.

5. Berry, J.D., Jones, S., Drebot, M.A., Andonov, A., Sabara, M., Yuan, X.Y., Weingartl, H., Fernando, L., Marszal, P., Gren, J., et al. (2004). Development and characterisation of neutralising monoclonal antibody to the SARS-coronavirus. J Virol Methods 120, 87–96. 10.1016/j.jviromet.2004.04.009.

6. Cameroni, E., Bowen, J.E., Rosen, L.E., Saliba, C., Zepeda, S.K., Culap, K., Pinto, D., VanBlargan, L.A., De Marco, A., di Iulio, J., et al. (2021). Broadly neutralizing antibodies overcome SARS-CoV-2 Omicron antigenic shift. Nature. 10.1038/s41586-021-04386-2.

7. Cerutti, G., Guo, Y., Zhou, T., Gorman, J., Lee, M., Rapp, M., Reddem, E.R., Yu, J., Bahna, F., Bimela, J., et al. (2021). Potent SARS-CoV-2 neutralizing antibodies directed against spike N-terminal domain target a single supersite. Cell Host Microbe. 10.1016/j.chom.2021.03.005.

8. Chaudhary, N., Weissman, D., and Whitehead, K.A. (2021). mRNA vaccines for infectious diseases: principles, delivery and clinical translation. Nat Rev Drug Discov 20, 817–838. 10.1038/s41573-021-00283-5.

9. Chi, X., Yan, R., Zhang, J., Zhang, G., Zhang, Y., Hao, M., Zhang, Z., Fan, P., Dong, Y., Yang, Y., et al. (2020). A potent neutralizing human antibody reveals the N-terminal domain of the Spike protein of SARS-CoV-2 as a site of vulnerability. 10.1101/2020.05.08.083964.

10. Coffman, R.L., Sher, A., and Seder, R.A. (2010). Vaccine adjuvants: putting innate immunity to work. Immunity 33, 492–503. 10.1016/j.immuni.2010.10.002.

11. Dai, L., Zheng, T., Xu, K., Han, Y., Xu, L., Huang, E., An, Y., Cheng, Y., Li, S., Liu, M., et al. (2020). A Universal Design of Betacoronavirus Vaccines against COVID-19, MERS, and SARS. Cell 182, 722–733 e711. 10.1016/j.cell.2020.06.035.

12. Fox, C.B., Kramer, R.M., Barnes, V.L., Dowling, Q.M., and Vedvick, T.S. (2013). Working together: interactions between vaccine antigens and adjuvants. Ther Adv Vaccines 1, 7–20. 10.1177/2051013613480144.

13. Fox, C.B., Orr, M.T., Van Hoeven, N., Parker, S.C., Mikasa, T.J., Phan, T., Beebe, E.A., Nana, G.I., Joshi, S.W., Tomai, M.A., et al. (2016). Adsorption of a synthetic TLR7/8 ligand to aluminum oxyhydroxide for enhanced vaccine adjuvant activity: A formulation approach. J Control Release 244, 98–107. 10.1016/j.jconrel.2016.11.011.

14. Francica, J.R., Flynn, B.J., Foulds, K.E., Noe, A.T., Werner, A.P., Moore, I.N., Gagne, M., Johnston, T.S., Tucker, C., Davis, R.L., et al. (2021). Protective antibodies elicited by SARS-CoV-2 spike protein vaccination are boosted in the lung after challenge in nonhuman primates. Sci Transl Med 13. 10.1126/scitranslmed.abi4547.

15. Freyn, A.W., Pine, M., Rosado, V.C., Benz, M., Muramatsu, H., Beattie, M., Tam, Y.K., Krammer, F., Palese, P., Nachbagauer, R., et al. (2021). Antigen modifications improve nucleoside-modified mRNA- based influenza virus vaccines in mice. Mol Ther Methods Clin Dev 22, 84–95. 10.1016/j.omtm.2021.06.003.

16. Freyn, A.W., Ramos da Silva, J., Rosado, V.C., Bliss, C.M., Pine, M., Mui, B.L., Tam, Y.K., Madden, T.D., de Souza Ferreira, L.C., Weissman, D., et al. (2020). A Multi-Targeting, Nucleoside-Modified mRNA Influenza Virus Vaccine Provides Broad Protection in Mice. Mol Ther 28, 1569–1584. 10.1016/j.ymthe.2020.04.018.

17. Gagne, M., Moliva, J.I., Foulds, K.E., Andrew, S.F., Flynn, B.J., Werner, A.P., Wagner, D.A., Teng, I.- T., Lin, B.C., Moore, C., et al. (2022). mRNA-1273 or mRNA-Omicron boost in vaccinated macaques elicits comparable B cell expansion, neutralizing antibodies and protection against Omicron. bioRxiv, 2022.2002.2003.479037. 10.1101/2022.02.03.479037.

18. Hastie, K.M., Li, H., Bedinger, D., Schendel, S.L., Dennison, S.M., Li, K., Rayaprolu, V., Yu, X., Mann, C., Zandonatti, M., et al. (2021). Defining variant-resistant epitopes targeted by SARS-CoV-2 antibodies: A global consortium study. Science 374, 472–478. 10.1126/science.abh2315.

19. Haynes, B.F., Corey, L., Fernandes, P., Gilbert, P.B., Hotez, P.J., Rao, S., Santos, M.R., Schuitemaker, H., Watson, M., and Arvin, A. (2020). Prospects for a safe COVID-19 vaccine. Sci Transl Med 12. 10.1126/scitranslmed.abe0948.

20. HogenEsch, H., O’Hagan, D.T., and Fox, C.B. (2018). Optimizing the utilization of aluminum adjuvants in vaccines: you might just get what you want. NPJ Vaccines 3, 51. 10.1038/s41541-018-0089-x.

21. Kasturi, S.P., Rasheed, M.A.U., Havenar-Daughton, C., Pham, M., Legere, T., Sher, Z.J., Kovalenkov, Y., Gumber, S., Huang, J.Y., Gottardo, R., et al. (2020). 3M-052, a synthetic TLR-7/8 agonist, induces durable HIV-1 envelope-specific plasma cells and humoral immunity in nonhuman primates. Sci Immunol 5. 10.1126/sciimmunol.abb1025.

22. Korber, B., Fischer, W.M., Gnanakaran, S., Yoon, H., Theiler, J., Abfalterer, W., Hengartner, N., Giorgi, E.E., Bhattacharya, T., Foley, B., et al. (2020). Tracking Changes in SARS-CoV-2 Spike: Evidence that D614G Increases Infectivity of the COVID-19 Virus. Cell 182, 812–827 e819. 10.1016/j.cell.2020.06.043.

23. Leist, S.R., Dinnon, K.H., 3rd, Schäfer, A., Tse, L.V., Okuda, K., Hou, Y.J., West, A., Edwards, C.E., Sanders, W., Fritch, E.J., et al. (2020). A Mouse-Adapted SARS-CoV-2 Induces Acute Lung Injury and Mortality in Standard Laboratory Mice. Cell 183, 1070–1085.e1012. 10.1016/j.cell.2020.09.050.

24. Levin, E.G., Lustig, Y., Cohen, C., Fluss, R., Indenbaum, V., Amit, S., Doolman, R., Asraf, K., Mendelson, E., Ziv, A., et al. (2021). Waning Immune Humoral Response to BNT162b2 Covid-19 Vaccine over 6 Months. N Engl J Med. 10.1056/NEJMoa2114583.

25. Li, D., Edwards, R.J., Manne, K., Martinez, D.R., Schafer, A., Alam, S.M., Wiehe, K., Lu, X., Parks, R., Sutherland, L.L., et al. (2021a). In vitro and in vivo functions of SARS-CoV-2 infection-enhancing and neutralizing antibodies. Cell 184, 4203–4219 e4232. 10.1016/j.cell.2021.06.021.

26. Li, D., Sempowski, G.D., Saunders, K.O., Acharya, P., and Haynes, B.F. (2021b). SARS-CoV-2 Neutralizing Antibodies for COVID-19 Prevention and Treatment. Annu Rev Med. 10.1146/annurev-med-042420-113838.

27. Maier, M.A., Jayaraman, M., Matsuda, S., Liu, J., Barros, S., Querbes, W., Tam, Y.K., Ansell, S.M., Kumar, V., Qin, J., et al. (2013). Biodegradable lipids enabling rapidly eliminated lipid nanoparticles for systemic delivery of RNAi therapeutics. Mol Ther 21, 1570–1578. 10.1038/mt.2013.124.

28. Marrack, P., McKee, A.S., and Munks, M.W. (2009). Towards an understanding of the adjuvant action of aluminium. Nat Rev Immunol 9, 287–293. 10.1038/nri2510.

29. Martinez, D.R., Schafer, A., Gobeil, S., Li, D., De la Cruz, G., Parks, R., Lu, X., Barr, M., Stalls, V., Janowska, K., et al. (2021a). A broadly cross-reactive antibody neutralizes and protects against sarbecovirus challenge in mice. Sci Transl Med, eabj7125. 10.1126/scitranslmed.abj7125.

30. Martinez, D.R., Schafer, A., Leist, S.R., De la Cruz, G., West, A., Atochina-Vasserman, E.N., Lindesmith, L.C., Pardi, N., Parks, R., Barr, M., et al. (2021b). Chimeric spike mRNA vaccines protect against Sarbecovirus challenge in mice. Science 373, 991–998. 10.1126/science.abi4506.

31. McCallum, M., De Marco, A., Lempp, F.A., Tortorici, M.A., Pinto, D., Walls, A.C., Beltramello, M., Chen, A., Liu, Z., Zatta, F., et al. (2021). N-terminal domain antigenic mapping reveals a site of vulnerability for SARS-CoV-2. Cell. 10.1016/j.cell.2021.03.028.

32. Menachery, V.D., Yount, B.L., Jr., Debbink, K., Agnihothram, S., Gralinski, L.E., Plante, J.A., Graham, R.L., Scobey, T., Ge, X.Y., Donaldson, E.F., et al. (2015). A SARS-like cluster of circulating bat coronaviruses shows potential for human emergence. Nat Med 21, 1508–1513. 10.1038/nm.3985.

33. Naldini, L., Blomer, U., Gage, F.H., Trono, D., and Verma, I.M. (1996). Efficient transfer, integration, and sustained long-term expression of the transgene in adult rat brains injected with a lentiviral vector. Proc Natl Acad Sci U S A 93, 11382–11388. 10.1073/pnas.93.21.11382.

34. Pardi, N., Hogan, M.J., Porter, F.W., and Weissman, D. (2018). mRNA vaccines - a new era in vaccinology. Nat Rev Drug Discov 17, 261–279. 10.1038/nrd.2017.243.

35. Pardi, N., Hogan, M.J., and Weissman, D. (2020). Recent advances in mRNA vaccine technology. Curr Opin Immunol 65, 14–20. 10.1016/j.coi.2020.01.008.

36. Pino, M., Abid, T., Pereira Ribeiro, S., Edara, V.V., Floyd, K., Smith, J.C., Latif, M.B., Pacheco-Sanchez, G., Dutta, D., Wang, S., et al. (2021). A yeast expressed RBD-based SARS-CoV-2 vaccine formulated with 3M-052-alum adjuvant promotes protective efficacy in non-human primates. Sci Immunol 6. 10.1126/sciimmunol.abh3634.

37. Pinto, D., Park, Y.J., Beltramello, M., Walls, A.C., Tortorici, M.A., Bianchi, S., Jaconi, S., Culap, K., Zatta, F., De Marco, A., et al. (2020). Cross-neutralization of SARS-CoV-2 by a human monoclonal SARS-CoV antibody. Nature 583, 290–295. 10.1038/s41586-020-2349-y.

38. Polack, F.P., Thomas, S.J., Kitchin, N., Absalon, J., Gurtman, A., Lockhart, S., Perez, J.L., Perez Marc, G., Moreira, E.D., Zerbini, C., et al. (2020). Safety and Efficacy of the BNT162b2 mRNA Covid-19 Vaccine. N Engl J Med 383, 2603–2615. 10.1056/NEJMoa2034577.

39. Roberts, A., Deming, D., Paddock, C.D., Cheng, A., Yount, B., Vogel, L., Herman, B.D., Sheahan, T., Heise, M., Genrich, G.L., et al. (2007). A mouse-adapted SARS-coronavirus causes disease and mortality in BALB/c mice. PLoS Pathog 3, e5. 10.1371/journal.ppat.0030005.

40. Routhu, N.K., Cheedarla, N., Bollimpelli, V.S., Gangadhara, S., Edara, V.V., Lai, L., Sahoo, A., Shiferaw, A., Styles, T.M., Floyd, K., et al. (2021). SARS-CoV-2 RBD trimer protein adjuvanted with Alum-3M-052 protects from SARS-CoV-2 infection and immune pathology in the lung. Nat Commun 12, 3587. 10.1038/s41467-021-23942-y.

41. Saunders, K.O., Lee, E., Parks, R., Martinez, D.R., Li, D., Chen, H., Edwards, R.J., Gobeil, S., Barr, M., Mansouri, K., et al. (2021). Neutralizing antibody vaccine for pandemic and pre-emergent coronaviruses. Nature 594, 553–559. 10.1038/s41586-021-03594-0.

42. Schmidt, F., Weisblum, Y., Rutkowska, M., Poston, D., DaSilva, J., Zhang, F., Bednarski, E., Cho, A., Schaefer-Babajew, D.J., Gaebler, C., et al. (2021). High genetic barrier to SARS-CoV-2 polyclonal neutralizing antibody escape. Nature 600, 512–516. 10.1038/s41586-021-04005-0.

43. Smirnov, D., Schmidt, J.J., Capecchi, J.T., and Wightman, P.D. (2011). Vaccine adjuvant activity of 3M- 052: an imidazoquinoline designed for local activity without systemic cytokine induction. Vaccine 29, 5434–5442. 10.1016/j.vaccine.2011.05.061.

44. Voysey, M., Clemens, S.A.C., Madhi, S.A., Weckx, L.Y., Folegatti, P.M., Aley, P.K., Angus, B., Baillie, V.L., Barnabas, S.L., Bhorat, Q.E., et al. (2021). Safety and efficacy of the ChAdOx1 nCoV-19 vaccine (AZD1222) against SARS-CoV-2: an interim analysis of four randomised controlled trials in Brazil, South Africa, and the UK. Lancet 397, 99–111. 10.1016/S0140-6736(20)32661-1.

45. Wang, L., and Cheng, G. (2021). Sequence analysis of the emerging SARS-CoV-2 variant Omicron in South Africa. J Med Virol. 10.1002/jmv.27516.

46. Wolfel, R., Corman, V.M., Guggemos, W., Seilmaier, M., Zange, S., Muller, M.A., Niemeyer, D., Jones, T.C., Vollmar, P., Rothe, C., et al. (2020). Virological assessment of hospitalized patients with COVID- 2019. Nature 581, 465–469. 10.1038/s41586-020-2196-x.

47. Wrapp, D., Wang, N., Corbett, K.S., Goldsmith, J.A., Hsieh, C.L., Abiona, O., Graham, B.S., and McLellan, J.S. (2020). Cryo-EM structure of the 2019-nCoV spike in the prefusion conformation. Science 367, 1260–1263. 10.1126/science.abb2507.

48. Yang, J., Wang, W., Chen, Z., Lu, S., Yang, F., Bi, Z., Bao, L., Mo, F., Li, X., Huang, Y., et al. (2020). A vaccine targeting the RBD of the S protein of SARS-CoV-2 induces protective immunity. Nature 586, 572–577. 10.1038/s41586-020-2599-8.

49. Yu, J., Tostanoski, L.H., Peter, L., Mercado, N.B., McMahan, K., Mahrokhian, S.H., Nkolola, J.P., Liu, J., Li, Z., Chandrashekar, A., et al. (2020). DNA vaccine protection against SARS-CoV-2 in rhesus macaques. Science 369, 806–811. 10.1126/science.abc6284.

50. Zhou, T., Teng, I.T., Olia, A.S., Cerutti, G., Gorman, J., Nazzari, A., Shi, W., Tsybovsky, Y., Wang, L., Wang, S., et al. (2020). Structure-Based Design with Tag-Based Purification and In-Process Biotinylation Enable Streamlined Development of SARS-CoV-2 Spike Molecular Probes. Cell Rep 33, 108322. 10.1016/j.celrep.2020.108322.

51. Zivanov, J., Nakane, T., Forsberg, B.O., Kimanius, D., Hagen, W.J., Lindahl, E., and Scheres, S.H. (2018). New tools for automated high-resolution cryo-EM structure determination in RELION-3. Elife 7. 10.7554/eLife.42166.

