## Supplementary Raw Data for Histopathologic Analysis for "Breadth of SARS-CoV-2 Neutralization and Protection Induced by a Nanoparticle Vaccine"

**1. Pathology data for the SARS-CoV-2 WA-1 challenge studies related to Figure 2.** Haematoxylin and eosin (H&E) staining histology and immunohistochemistry (IHC) staining of lung tissue. Each column shows results from an individual macaque. The macaque identification number is shown above each column. Red arrows indicate site of antigen positivity. All images are shown at 10× magnification.

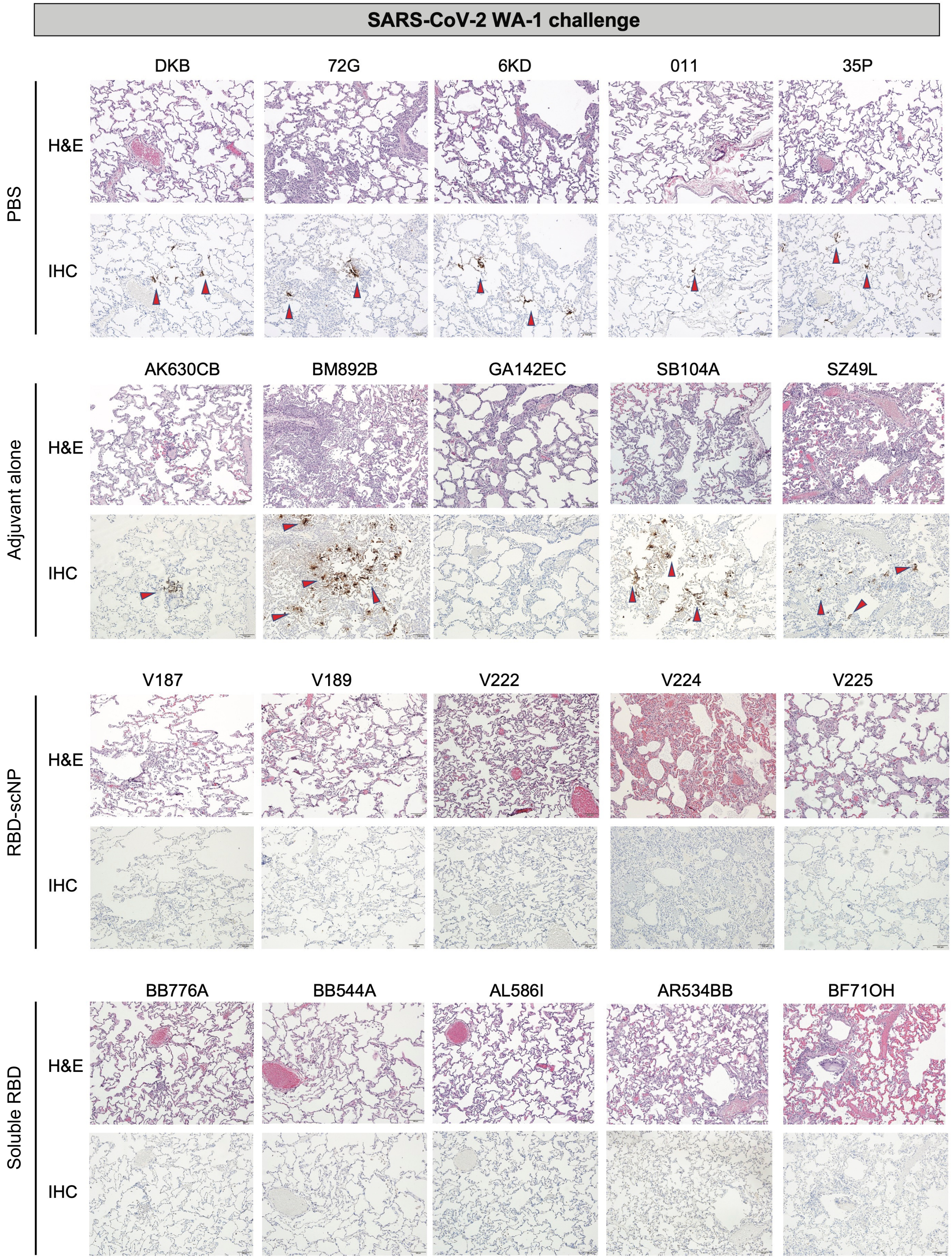

**2. Pathology data for the SARS-CoV-2 beta challenge studies related to Figure 2.** Haematoxylin and eosin (H&E) staining histology and immunohistochemistry (IHC) staining of lung tissue. Each column shows results from an individual macaque. The macaque identification number is shown above each column. Red arrows indicate site of antigen positivity. All images are shown at 10× magnification.

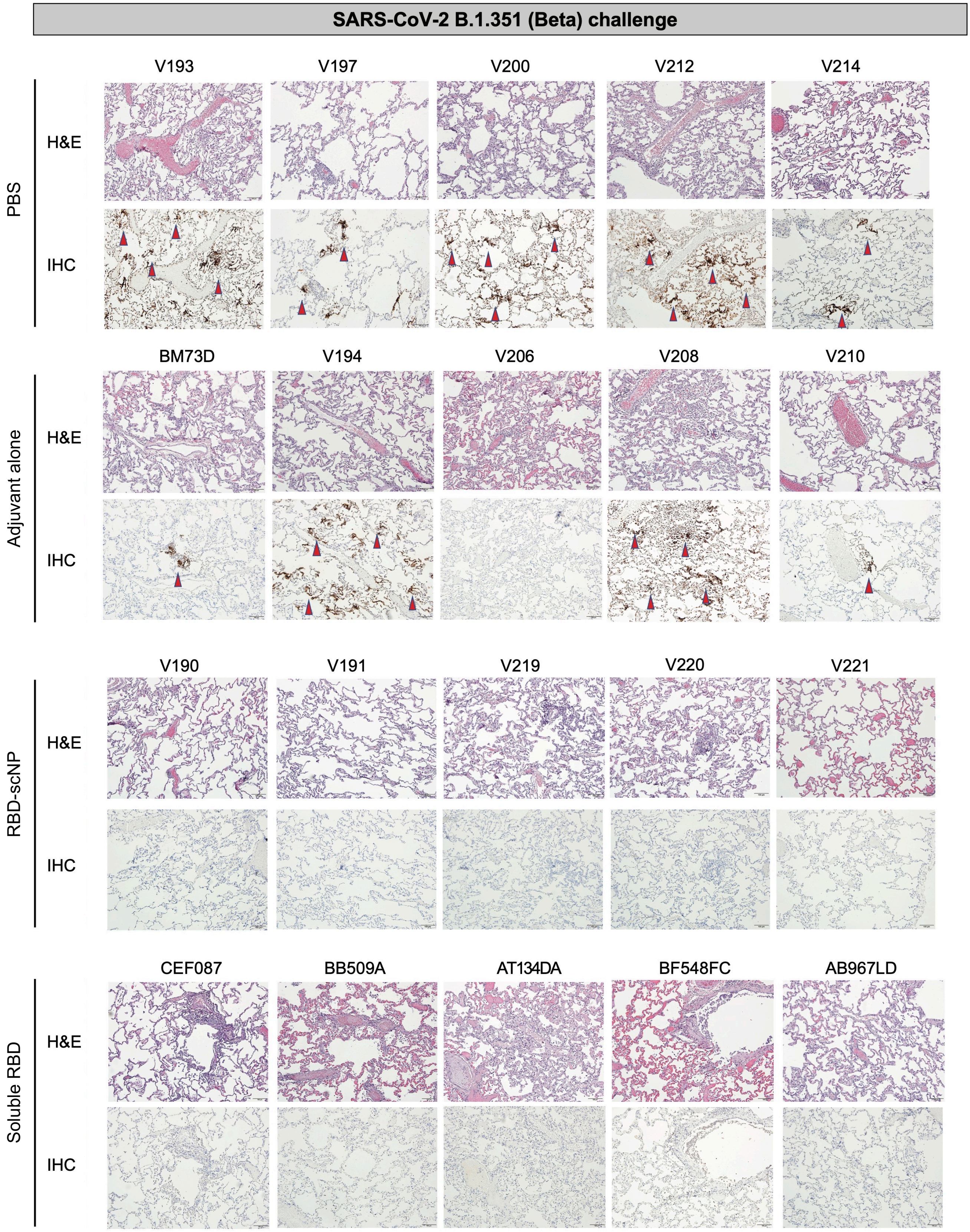

**3. Pathology data for the SARS-CoV-2 WA-1 challenge and beta challenge studies related to Figure 2.** Scoring of H&E staining and IHC staining of macaque lung tissue. H&E staining (inflammation): −, minimal to absent; +/−, minimal to mild; +, mild to moderate; ++, moderate to severe; +++, severe. IHC staining (SARS-CoV-2 nucleocapsid antigen-positive foci): −, no SARS-CoV-2 antigen detected; +/−, rare or occasional; +, occasional or multiple; ++, multiple or numerous (and foci often larger); +++, numerous. Lung region abbreviation: left caudal (Lc), right middle (Rm), and right caudal (Rc).

| | WA-1 challenge | | | B.1.351 (beta) challenge | | |
|---|---|---|---|---|---|---|
| Group | Animal ID | H&E (Lc; Rm; Rc) | COVID IHC (Lc; Rm; Rc) | Animal ID | H&E (Lc; Rm; Rc) | COVID IHC (Lc; Rm; Rc) |
| PBS | DKB | +/-; +/-; +/- | +; ++; ++ | V193 | ++; +; + | +++; +; +++ |
| PBS | 726 | ++; +; ++ | +++; +++; +++ | V197 | +/-; +; ++ | ++; ++; ++ |
| PBS | 6KD | +/-; +; + | -; ++; +++ | V200 | ++; +; + | +++; +++; ++ |
| PBS | 11 | +; +/-; +/- | +/-; +/-; + | V212 | +; ++; ++ | +++; ++; +++ |
| PBS | 35P | +; +; ++ | ++; ++; +++ | V214 | +; +/-; +/- | +; +; +/- |
| Adjuvant alone | AK630CB | +/-; +/-; + | +/-; +/-; + | BM73D | ++; ++; + | -; +; +/- |
| Adjuvant alone | BM892B | ++; +; + | +++; +; ++ | V194 | +/-; +/-; + | +++; -; ++ |
| Adjuvant alone | GA142EC | +; +; +/- | -; -; - | V206 | +; ++; ++ | -; -; - |
| Adjuvant alone | SB104A | +; +/-; ++ | +/-; +/-; ++ | V208 | ++; +/-; ++ | +++; +++; ++ |
| Adjuvant alone | SZ49L | +/-; +/-; + | +; +/-; + | V210 | +/-; +; + | +/-; +/-; +/- |
| RBD-scNP | V187 | +; +/-; ++ | -; -; - | V190 | ++; +/-; +/- | -; -; - |
| RBD-scNP | V189 | +; +; +/- | -; -; - | V191 | +; +/-; +/- | -; -; - |
| RBD-scNP | V222 | +/-; +; +/- | -; -; - | V219 | ++; +; + | -; -; - |
| RBD-scNP | V224 | ++; ++; + | -; -; - | V220 | ++; +/-; + | -; -; - |
| RBD-scNP | V225 | +/-; +; + | -; -; - | V221 | +/-; +/-; + | -; -; - |
| soluble RBD | BB776A | +/-; +/-; +/- | -; -; - | CEF087 | +; +; + | -; -; - |
| soluble RBD | BB544A | +/-; +/-; +/- | -; -; - | BB509A | +; +; +/- | -; -; - |
| soluble RBD | AL586I | +/-; +/-; +/- | -; -; - | AT134DA | ++; ++; + | +; -; - |
| soluble RBD | AR534BB | +; +; + | -; -; - | BF548FC | +; +; + | +/-; -; - |
| soluble RBD | BF71OH | ++; ++; + | -; -; - | AB967LD | +; +/-; ++ | -; -; - |

**4. Haematoxylin and eosin (H&E) staining histology and immunohistochemistry (IHC) staining of lung tissue collected four days after SARS-CoV-2 WA-1 intratracheal and intranasal challenge.** Each column shows results from an individual macaque. The macaque identification number is shown above each column. Red arrows indicate site of antigen positivity. All images are shown at 10× magnification. **(Part 1) Related to Figure 3.**

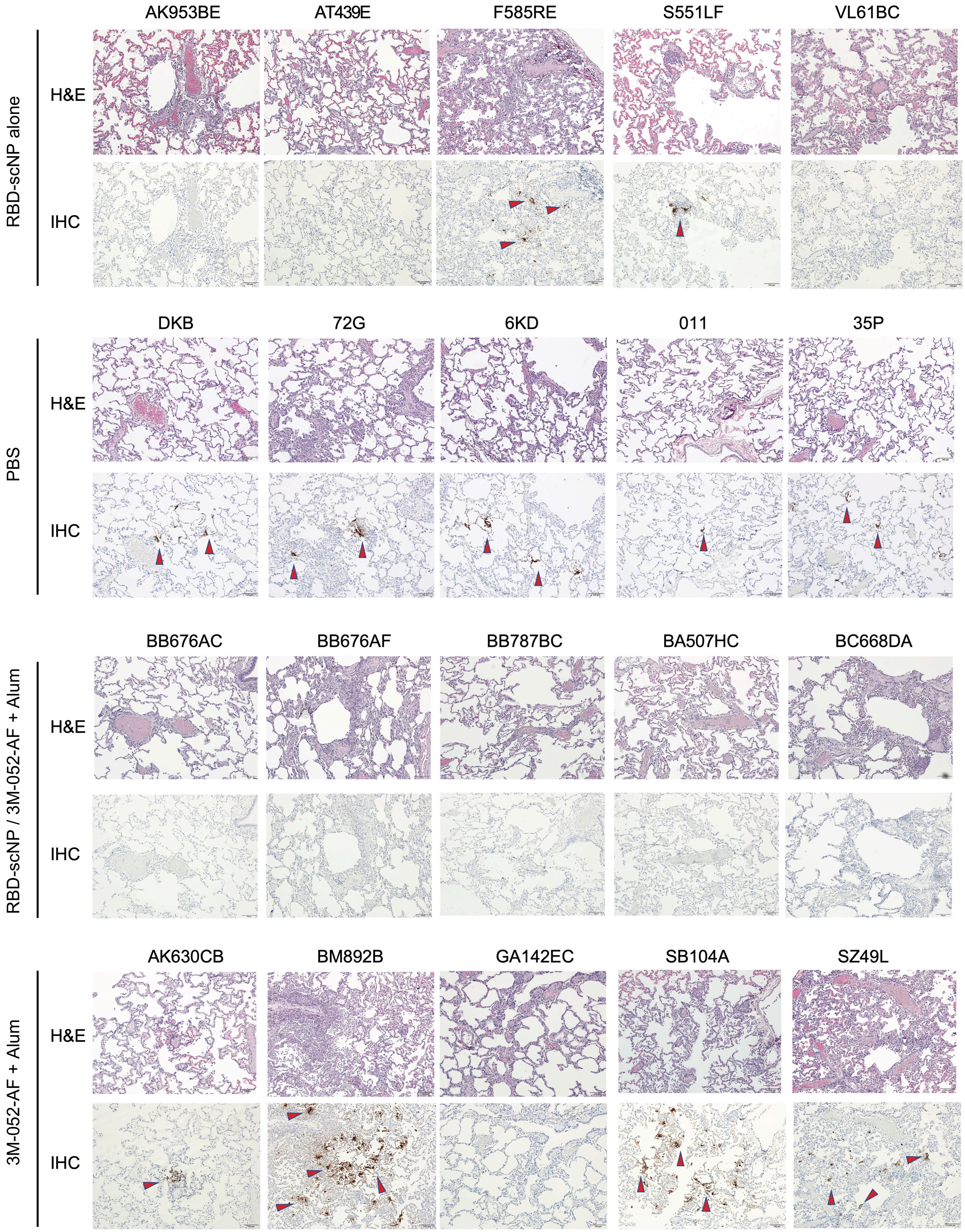

**5. Haematoxylin and eosin (H&E) staining histology and immunohistochemistry (IHC) staining of lung tissue collected four days after SARS-CoV-2 WA-1 intratracheal and intranasal challenge.** Each column shows results from an individual macaque. The macaque identification number is shown above each column. Red arrows indicate site of antigen positivity. All images are shown at 10× magnification. **(Part 2) Related to Figure 3.**

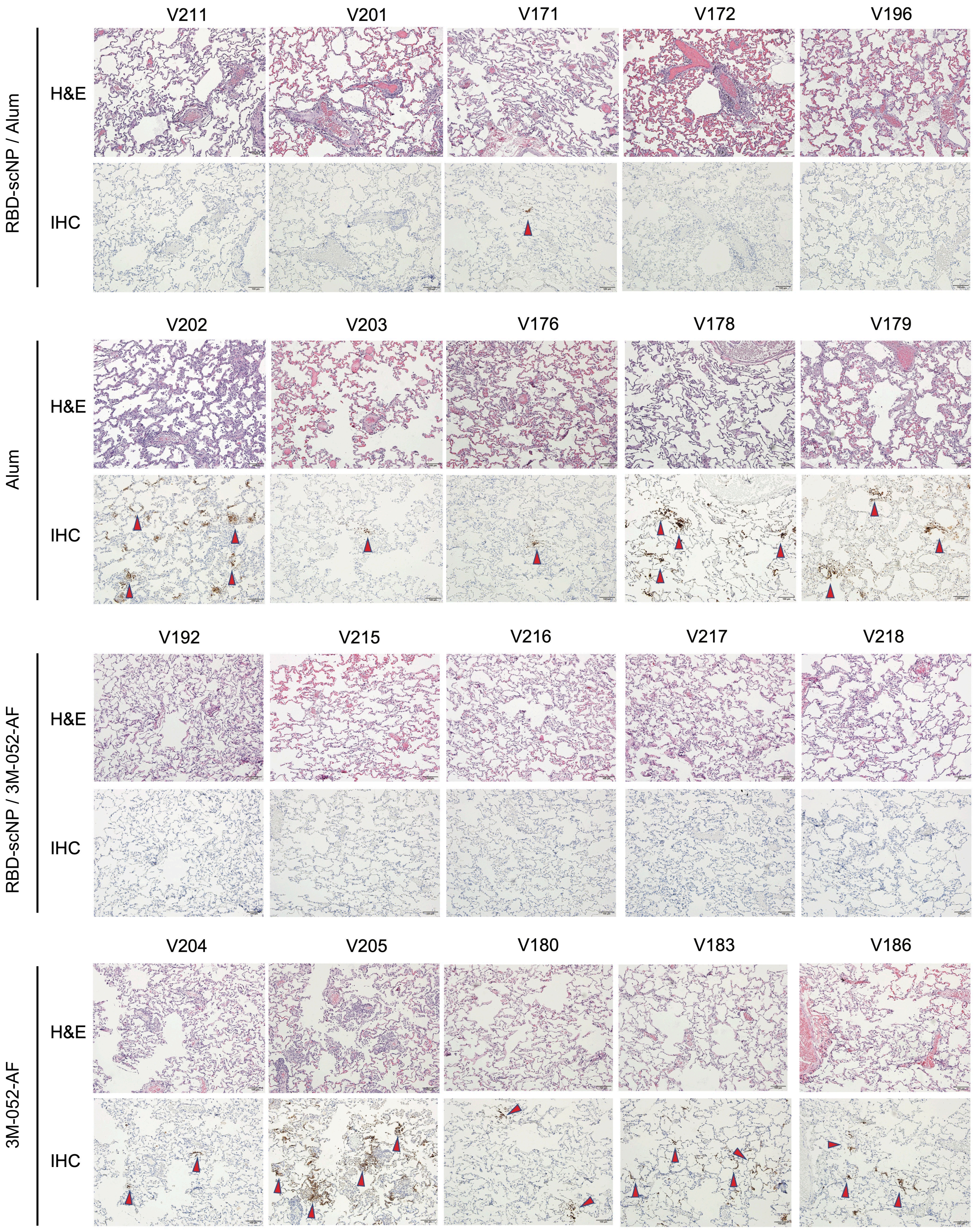

**6. Scoring of haematoxylin and eosin (H&E) staining and immunohistochemistry (IHC) staining of macaque lung tissue collected four days after challenge.** H&E staining (inflammation): −, minimal to absent; +/−, minimal to mild; +, mild to moderate; ++, moderate to severe; +++, severe. IHC staining (SARS-CoV-2 nucleocapsid antigen-positive foci): −, no SARS-CoV-2 antigen detected; +/−, rare or occasional; +, occasional or multiple; ++, multiple or numerous (and foci often larger); +++, numerous. Lung region abbreviation: left caudal (Lc), right middle (Rm), and right caudal (Rc). **Related to Figure 3.**

| Group | Animal ID | H&E (Lc; Rm; Rc) | IHC (Lc; Rm; Rc) |
|---|---|---|---|
| RBD-scNP alone | AK953BE | +/-; +/-; ++ | -; -; - |
| | AT439E | +; +; ++ | -; -; - |
| | F585RE | +/-; +/-; ++ | +; -; - |
| | S551LF | +; ++; + | +/-; -; - |
| | VL61BC | +; +; + | -; -; - |
| PBS | DKB | +/-; +/-; +/- | +; ++; ++ |
| | 726 | ++; +; ++ | +++; +++; +++ |
| | 6KD | +/-; +; + | -; ++; +++ |
| | 011 | +; +/-; +/- | +/-; +/-; + |
| | 35P | +; +; ++ | ++; ++; +++ |
| RBD-scNP / 3M-052-AF + Alum | BB676AC | +/-; +; + | -; -; - |
| | BB676AF | ++; ++; + | -; -; - |
| | BB787BC | ++; +; + | -; -; - |
| | BA507HC | +; +; +/- | -; -; - |
| | BC668DA | +/-; ++; ++ | -; -; - |
| 3M-052-AF + Alum | AK630CB | +; +; ++ | +/-; +/-; + |
| | BM892B | +++; +; ++ | +++; +; ++ |
| | GA142EC | +/-; +; + | -; -; - |
| | SB104A | +; +/-; ++ | +/-; +/-; ++ |
| | SZ49L | +/-; +; ++ | +; +/-; + |
| RBD-scNP / Alum | V211 | +/-; +/-; + | -; -; - |
| | V201 | +; ++; ++ | -; -; - |
| | V171 | +; +; +/- | +/-; -; +/- |
| | V172 | +; ++; ++ | -; -; - |
| | V196 | +; +; + | -; -; - |
| Alum | V202 | ++; +++; +++ | +++; +++; ++ |
| | V203 | +; +; + | +/-; +/-; +/- |
| | V176 | +/-; +; + | +/-; +; +/- |
| | V178 | +/-; +/-; + | +++; +; +++ |
| | V179 | +++; ++; ++ | ++; ++; ++ |
| RBD-scNP / 3M-052-AF | V192 | +/-; +/-; +/- | -; -; - |
| | V215 | +/-; +/-; + | -; -; - |
| | V216 | +/-; +/-; +/- | -; -; - |
| | V217 | +/-; +/-; + | -; -; - |
| | V218 | +/-; +; + | -; -; - |
| 3M-052-AF | V204 | +; +/-; + | +; +; + |
| | V205 | +; ++; ++ | +++; +++; +++ |
| | V180 | +; +; + | +; -; + |
| | V183 | ++; +; + | ++; +++; ++ |
| | V186 | +/-; +; + | ++; ++; ++ |

**7. Pathology data for the SARS-CoV-2 WA-1 challenge study related to Figure 4a-i.** Haematoxylin and eosin (H&E) staining histology and immunohistochemistry (IHC) staining of lung tissue. Each column shows results from an individual macaque. The macaque identification number is shown above each column. Red arrows indicate site of antigen positivity. All images are shown at 10× magnification.

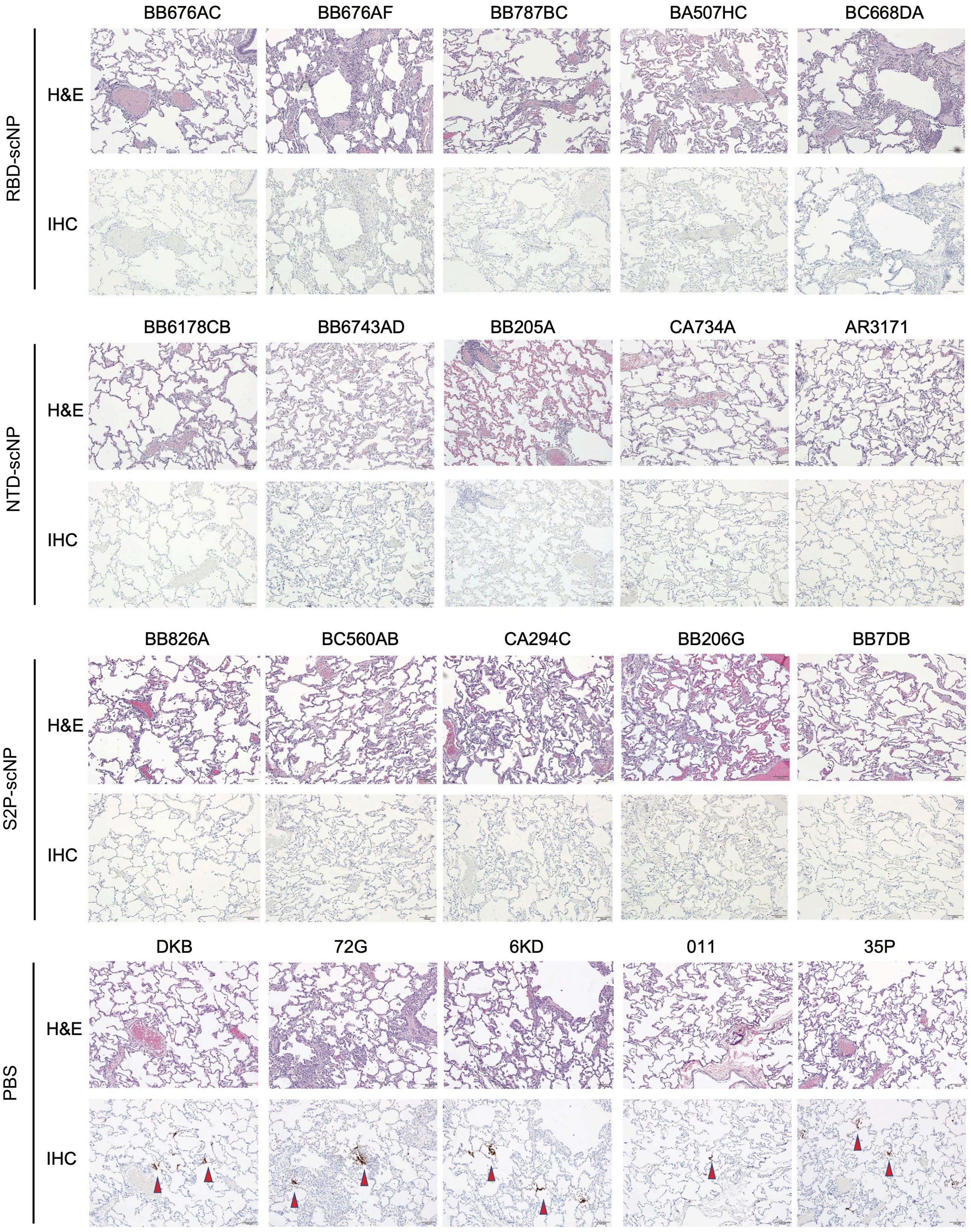

**8. Pathology data for the SARS-CoV-2 WA-1 challenge study related to Figure 4a-i.** Scoring of H&E staining and IHC staining of macaque lung tissue. H&E staining (inflammation): −, minimal to absent; +/−, minimal to mild; +, mild to moderate; ++, moderate to severe; +++, severe. IHC staining (SARS-CoV-2 nucleocapsid antigen-positive foci): −, no SARS-CoV-2 antigen detected; +/−, rare or occasional; +, occasional or multiple; ++, multiple or numerous (and foci often larger); +++, numerous. Lung region abbreviation: left caudal (Lc), right middle (Rm), and right caudal (Rc).

| **Group** | **Animal ID** | **H&E (Lc; Rm; Rc)** | **COVID IHC (Lc; Rm; Rc)** |
|---|---|---|---|
| RBD-scNP | BB676AC | +/-; +; + | -; -; - |
| RBD-scNP | BB676AF | ++; ++; + | -; -; - |
| RBD-scNP | BB787BC | ++; +; + | -; -; - |
| RBD-scNP | BA507HC | +; +; +/- | -; -; - |
| RBD-scNP | BC668DA | +/-; ++; ++ | -; -; - |
| NTD-scNP | BB6178CB | +/-; +; +/- | -; -; - |
| NTD-scNP | BB6743AD | +/-; +; + | -; -; - |
| NTD-scNP | BB205A | +/-; +/-; + | -; -; - |
| NTD-scNP | CA734A | +/-; +/-; +/- | -; -; - |
| NTD-scNP | AR3171 | +/-; +/-; +/- | -; -; - |
| S2P-scNP | BB826A | +/-; +; + | -; -; - |
| S2P-scNP | BC560AB | +; +/-; + | -; -; - |
| S2P-scNP | CA294C | ++*; +/-; +* | -; -; - |
| S2P-scNP | BB206G | +; +; + | -; -; - |
| S2P-scNP | BB7DB | +/-; +/-; +/- | -; -; - |
| PBS | DKB | +/-; +/-; +/- | +; ++; ++ |
| PBS | 726 | ++; +; ++ | +++; +++; +++ |
| PBS | 6KD | +/-; +; + | -; ++; +++ |
| PBS | 11 | +; +/-; +/- | +/-; +/-; + |
| PBS | 35P | +; +; ++ | ++; ++; +++ |

**9. Pathology data for the SARS-CoV-2 WA-1 challenge study related to Figure 4j-o.** Haematoxylin and eosin (H&E) staining histology and immunohistochemistry (IHC) staining of lung tissue. Each column shows results from an individual macaque. The macaque identification number is shown above each column. Red arrows indicate site of antigen positivity. All images are shown at 10× magnification.

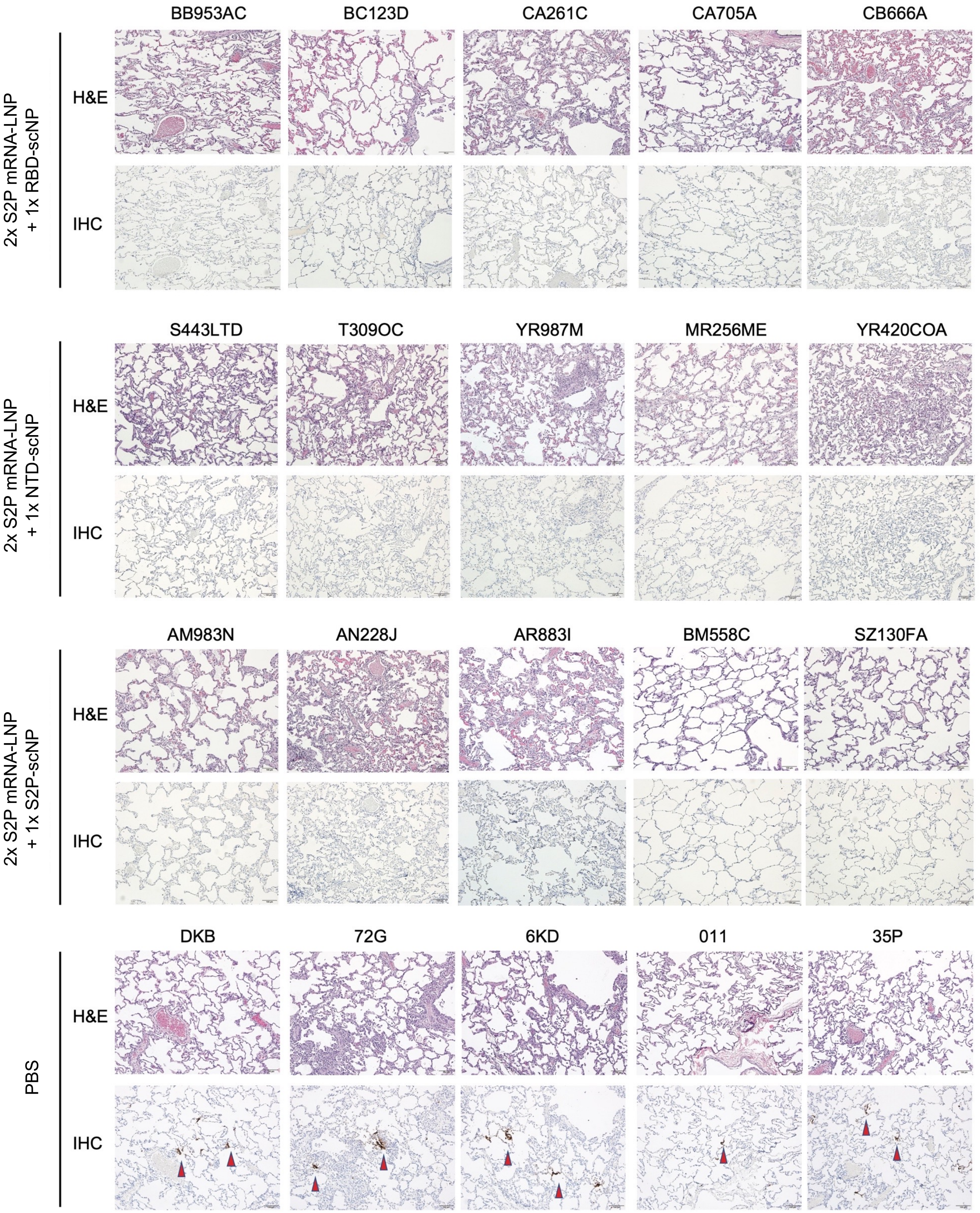

**10. Pathology data for the SARS-CoV-2 WA-1 challenge study related to Figure 4j-o.** Scoring of H&E staining and IHC staining of macaque lung tissue. H&E staining (inflammation): −, minimal to absent; +/−, minimal to mild; +, mild to moderate; ++, moderate to severe; +++, severe. IHC staining (SARS-CoV-2 nucleocapsid antigen-positive foci): −, no SARS-CoV-2 antigen detected; +/−, rare or occasional; +, occasional or multiple; ++, multiple or numerous (and foci often larger); +++, numerous. Lung region abbreviation: left caudal (Lc), right middle (Rm), and right caudal (Rc).

| Group | Animal ID | H&E (Lc; Rm; Rc) | COVID IHC (Lc; Rm; Rc) |
|---|---|---|---|
| 2x S2P mRNA-LNP + 1x RBD-scNP | BB953AC | +/-; +/-; +/- | -; -; - |
| 2x S2P mRNA-LNP + 1x RBD-scNP | BC123D | +/-; -; +/- | -; -; - |
| 2x S2P mRNA-LNP + 1x RBD-scNP | CA261C | +/-; +; + | -; -; - |
| 2x S2P mRNA-LNP + 1x RBD-scNP | CA705A | +/-; -; - | -; -; - |
| 2x S2P mRNA-LNP + 1x RBD-scNP | CB666A | +/-; +; + | -; -; - |
| 2x S2P mRNA-LNP + 1x NTD-scNP | S443LTD | +/-; +; ++ | -; -; - |
| 2x S2P mRNA-LNP + 1x NTD-scNP | T309OC | +; +/-; + | -; -; - |
| 2x S2P mRNA-LNP + 1x NTD-scNP | YR987M | +; +; ++ | -; -; - |
| 2x S2P mRNA-LNP + 1x NTD-scNP | MR256ME | +/-; +++; + | -; -; - |
| 2x S2P mRNA-LNP + 1x NTD-scNP | YR420COA | +; ++; ++ | -; -; - |
| 2x S2P mRNA-LNP + 1x S2P-scNP | AM983N | +; ++; +/- | -; -; - |
| 2x S2P mRNA-LNP + 1x S2P-scNP | AN228J | +; ++; ++ | -; -; - |
| 2x S2P mRNA-LNP + 1x S2P-scNP | AR883I | +; ++; + | -; -; - |
| 2x S2P mRNA-LNP + 1x S2P-scNP | BM558C | +/-; +; +/- | -; -; - |
| 2x S2P mRNA-LNP + 1x S2P-scNP | SZ130FA | +/-; +/-; + | -; -; - |
| PBS | DKB | +/-; +/-; +/- | +; ++; ++ |
| PBS | 726 | ++; +; ++ | +++; +++; +++ |
| PBS | 6KD | +/-; +; + | -; ++; +++ |
| PBS | 11 | +; +/-; +/- | +/-; +/-; + |
| PBS | 35P | +; +; ++ | ++; ++; +++ |
